# A proteomic atlas of organelle remodeling identifies lysosomal SNX3 as a regulator of Notch signaling in epidermal differentiation

**DOI:** 10.64898/2026.05.20.726558

**Authors:** Alex Hoover, Xinlei Sheng, Jinjun Gao, Jimmy Lee, Han Liu, Ben Taubman, Suman Suman, Shao-Yu Chen, Yingming Zhao, Xiaoyang Wu

**Author notes:** To whom correspondence should be addressed, Xiaoyang Wu, The University of Chicago, GCIS W408B, 929 E 57^th^ Street, Chicago, IL 60637, USA. These authors contributed equally (Alex Hoover, Xinlei Sheng).

## Abstract

Differentiation of epidermal keratinocytes is accompanied by profound reorganization of intracellular architecture, but how organelle remodeling interfaces with cell fate control is not well understood. Here, we generate a compartment-resolved proteomic map of keratinocyte differentiation and identify extensive remodeling of lysosomes, mitochondria, autophagic vesicles, plasma membrane, and nucleus. Differentiating keratinocytes display coordinated enrichment of lysosomal degradative machinery, vesicular trafficking factors, and mitochondrial metabolic proteins, revealing organelle remodeling as a prominent feature of epidermal differentiation. From the lysosomal proteome, we identify Sorting Nexin 3 (SNX3) as a critical regulator of epidermal homeostasis. SNX3 increasingly localizes to LAMP1-positive vesicles during differentiation, and its loss impairs epidermal differentiation, suppresses Notch signaling, and promotes proliferative gene expression. *In vivo*, *SNX3*-deficient skin grafts fail to maintain normal epidermal architecture and instead develop into squamous cell carcinoma. Mechanistically, SNX3 mediates efficient Notch receptor activation, as *SNX3* loss reduces nuclear Notch1 and NICD production, whereas NICD re-expression can rescue the differentiation defect. Our study defines a proteomic framework for organelle remodeling during epidermal differentiation and identifies lysosome-associated SNX3 as a key link between endolysosomal trafficking, Notch signaling, and epidermal tissue homeostasis.

## INTRODUCTION

Mammalian skin serves as the first line of defense against environmental insults. These essential barrier functions are sustained through continuous epidermal renewal, which is driven by epidermal stem/progenitor cells residing in the basal layer of the epidermis. Upon commitment to differentiation, these cells exit the cell cycle, detach from the basement membrane, and undergo a highly ordered terminal differentiation program as they migrate toward the skin surface^1^. This process culminates in the formation of enucleated squames encased within a rigid cornified envelope, establishing the physical and biochemical barrier of the epidermis^2,3^. Disruption of epidermal differentiation contributes to various skin diseases, including inflammatory skin disorders and skin cancers, underscoring the importance of understanding the molecular mechanisms governing this process^4^.

Epidermal differentiation is accompanied by profound transcriptional changes and extensive remodeling of intracellular architecture^5–8^. As keratinocytes progress through suprabasal layers, they undergo dramatic changes in subcellular compartments^2,9,10^., including mitochondrial remodeling, alterations in plasma membrane composition, activation of autophagy pathways, and ultimately the degradation of all intracellular organelles and the nucleus^8,11^. These events are tightly coordinated with differentiation-associated signaling pathways, among which Notch signaling plays a central role in enforcing cell fate commitment and suppressing keratinocyte proliferation^12–14^. While the transcriptional regulation and signal transduction pathways controlling epidermal differentiation have been extensively studied, how differentiation-associated signaling is coordinated with large-scale remodeling of subcellular organelles remains poorly understood.

One major challenge in addressing this question is that changes in organelle composition and protein localization are not necessarily accompanied by corresponding changes in keratinocyte transcriptome. Dynamic trafficking of proteins to and from specific organelles can profoundly influence signaling output, metabolism, and cell fate decisions^15–17^. Consequently, transcriptomic approaches alone are insufficient to capture these regulatory events. Advances in quantitative proteomics now enable systematic interrogation of protein localization across subcellular compartments, providing a powerful approach to define organelle-specific proteome remodeling in a state-dependent manner. Here, we employed a quantitative proteomics strategy to generate an organelle-resolved atlas of proteome remodeling during keratinocyte differentiation. By profiling multiple subcellular compartments, including lysosomes, autophagosomes, mitochondria, plasma membrane, and nucleus, we uncovered widespread and coordinated changes in organelle-associated proteins as keratinocytes differentiate. Among these, we identified Sorting Nexin 3 (SNX3) as a lysosome-associated protein that becomes enriched during differentiation. Functional studies revealed that loss of *SNX3* impairs epidermal differentiation, reduces Notch signaling activity, and diminishes nuclear accumulation of the Notch intracellular domain (NICD). Furthermore, SNX3 deficiency predisposes epidermal cells to tumorigenesis, highlighting its critical role in maintaining skin tissue homeostasis. Together, our findings establish organelle proteome remodeling as a fundamental feature of epidermal differentiation and identify lysosomal SNX3 as a key regulator linking endolysosomal trafficking to Notch-dependent cell fate control in the skin.

## RESULTS

### Keratinocytes undergo extensive subcellular remodeling during epidermal differentiation

To characterize subcellular remodeling upon epidermal differentiation, we performed live-cell imaging of lysosomes, mitochondria, and endoplasmic reticulum (ER) in undifferentiated and differentiated keratinocytes *in vitro*. In proliferating basal keratinocytes, lysosomes exhibited a punctate morphology and were largely confined to the perinuclear region. By contrast, upon calcium-induced differentiation, lysosomes underwent striking morphological changes, characterized by the appearance of elongated, tubular structures (Fig. 1A). Lysosome tubulation has previously been associated with increased autophagic flux and autolysosome reformation, consistent with the notion that autophagy-related pathways are activated early during keratinocyte differentiation^18,19^. Mitochondrial morphology also exhibited marked changes during differentiation. Undifferentiated keratinocytes displayed an interconnected, tubular mitochondrial network, whereas differentiated cells had a more fragmented mitochondrial architecture (Fig. 1B), a pattern that has been reported in three-dimensional (3D) human skin organoid models^11^. In addition, ER morphology was altered during differentiation. While undifferentiated keratinocytes maintained an extended and continuous ER network, differentiated cells frequently showed fragmented and discontinuous ER structures (Fig. 1C), consistent with increased ER stress during epidermal differentiation^20,21^.

**Figure 1.**
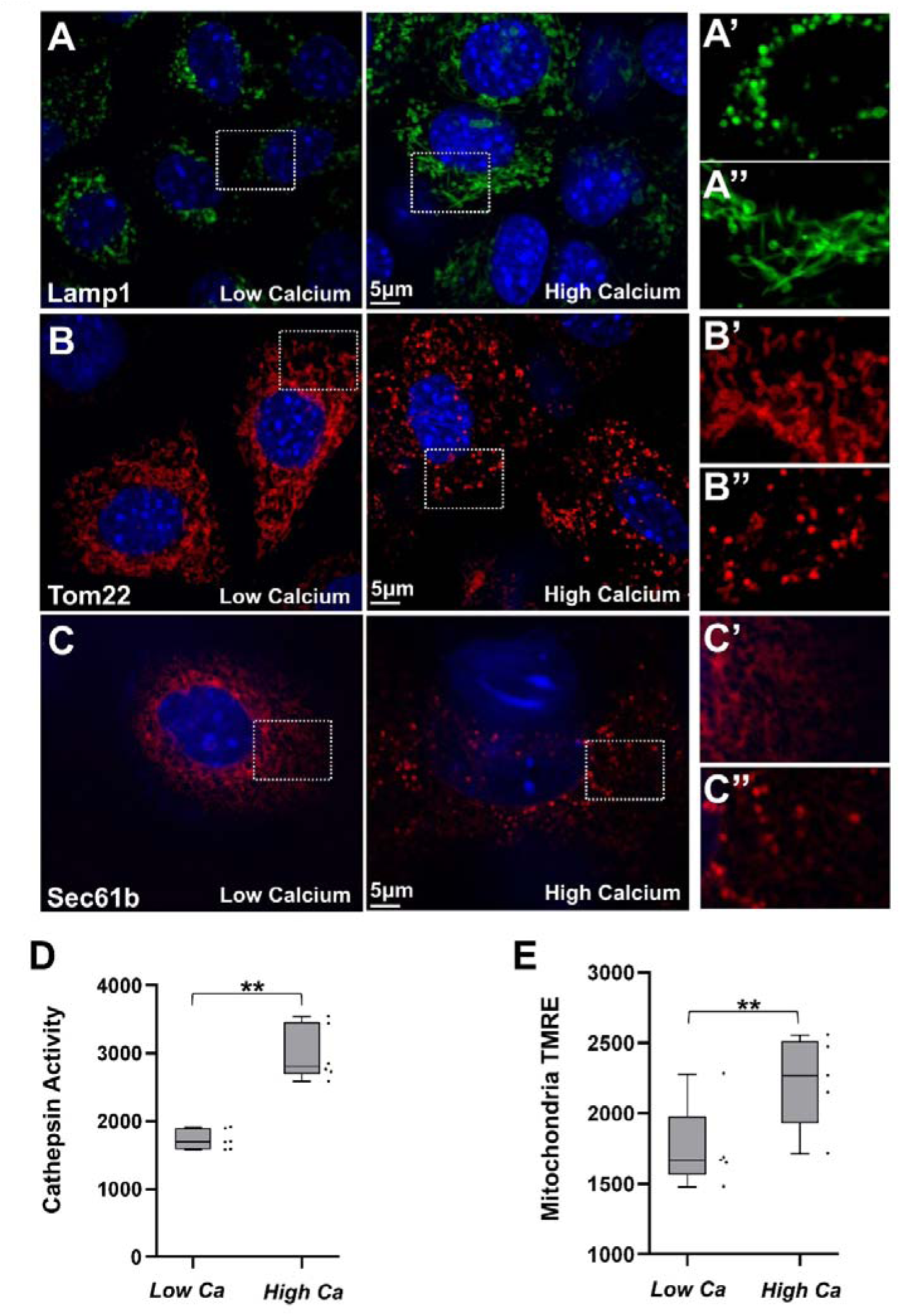
Keratinocytes Undergo Subcellular and Metabolic Changes During Differentiation. (A-C) Super resolution confocal images of organelle morphology changes during keratinocyte differentiation. Lysosome, mitochondria, and ER were tagged with Lamp1, Tom22, and Sec61b attached to fluorescent proteins. Lysosome tubular morphology becomes apparent during differentiation. Mitochondria and ER both exhibit a fragmentation phenomenon during differentiation. **(D)** Protein lysates from undifferentiated and differentiated keratinocytes were used to measure Cathepsin D activity. Differentiated cell lysates exhibited higher cathepsin D activity. N=6; **: P<0.05 (Student’s t-test). **(E)** Mitochondrial activity was measured via tetramethylrhodamine ethyl ester (TMRE) to monitor membrane potential in undifferentiated and differentiated cells. Differentiated cells showed higher TMRE fluorescence. N=5; P<0.05 (Student’s t-test).

To functionally assess lysosomal activity, we measured cathepsin D enzymatic activity in undifferentiated and differentiated keratinocytes. Differentiated cells exhibited significantly higher cathepsin D activity compared to undifferentiated controls (Fig. 1D), supporting enhanced lysosomal degradative capacity upon epidermal differentiation. To monitor remodeling of mitochondria, we quantified tetramethylrhodamine ethyl ester (TMRE) fluorescence in undifferentiated and differentiated keratinocytes to assess mitochondrial membrane potential. Differentiated cells showed increased TMRE signal intensity (Fig. 1E), suggesting elevated membrane potential and a shift in mitochondrial functional state upon differentiation. Cellular metabolism, especially lysosome function has been suggested to be necessary for skin differentiation^22^. We used bafilomycinA to inhibit lysosomal acidification and found that BafA inhibited calcium switch-induced differentiation of primary mouse keratinocytes, as determined by immunoblots of *loricrin* expression (Supplementary Fig. 1A-B).

### Quantitative characterization of subcellular compartment-specific proteomics during epidermal differentiation

Dynamic redistribution of proteins among subcellular compartments can drive critical signaling and metabolic events during cell fate transitions, yet such changes are often uncoupled from alterations in total gene expression. As keratinocytes undergo extensive morphological and functional remodeling during differentiation, we sought to systematically characterize proteome changes across distinct subcellular compartments. To this end, we performed quantitative SILAC (Stable Isotope Labeling by Amino acids in Cell culture)-based mass spectrometry on enriched subcellular fractions isolated from undifferentiated, actively proliferating keratinocytes and keratinocytes induced to differentiate by calcium switch. To enrich specific subcellular compartments, we first employed an organelle immunoprecipitation strategy that enables rapid and selective isolation of subcellular structures^23–25^. Lysosomes, mitochondria, and LC3-positive autophagic vesicles were labeled by expressing *LAMP1*, *TOM22*, or *LC3* fused to a twin-strep affinity tag, followed by streptavidin-mediated purification after mechanical cell lysis. This approach provides efficient enrichment of intact organelles and avoids the extended processing times required for traditional ultracentrifugation-based fractionation. Correct targeting of LAMP1 and TOM22 with twin-strep tag was confirmed by robust colocalization with LysoTracker and MitoTracker respectively (Fig. 2A), as well as by immunoblot analysis demonstrating strong enrichment of compartment-specific markers following isolation by twin-strep affinity tag (Supplementary Fig. 2A-B).

**Figure 2.**
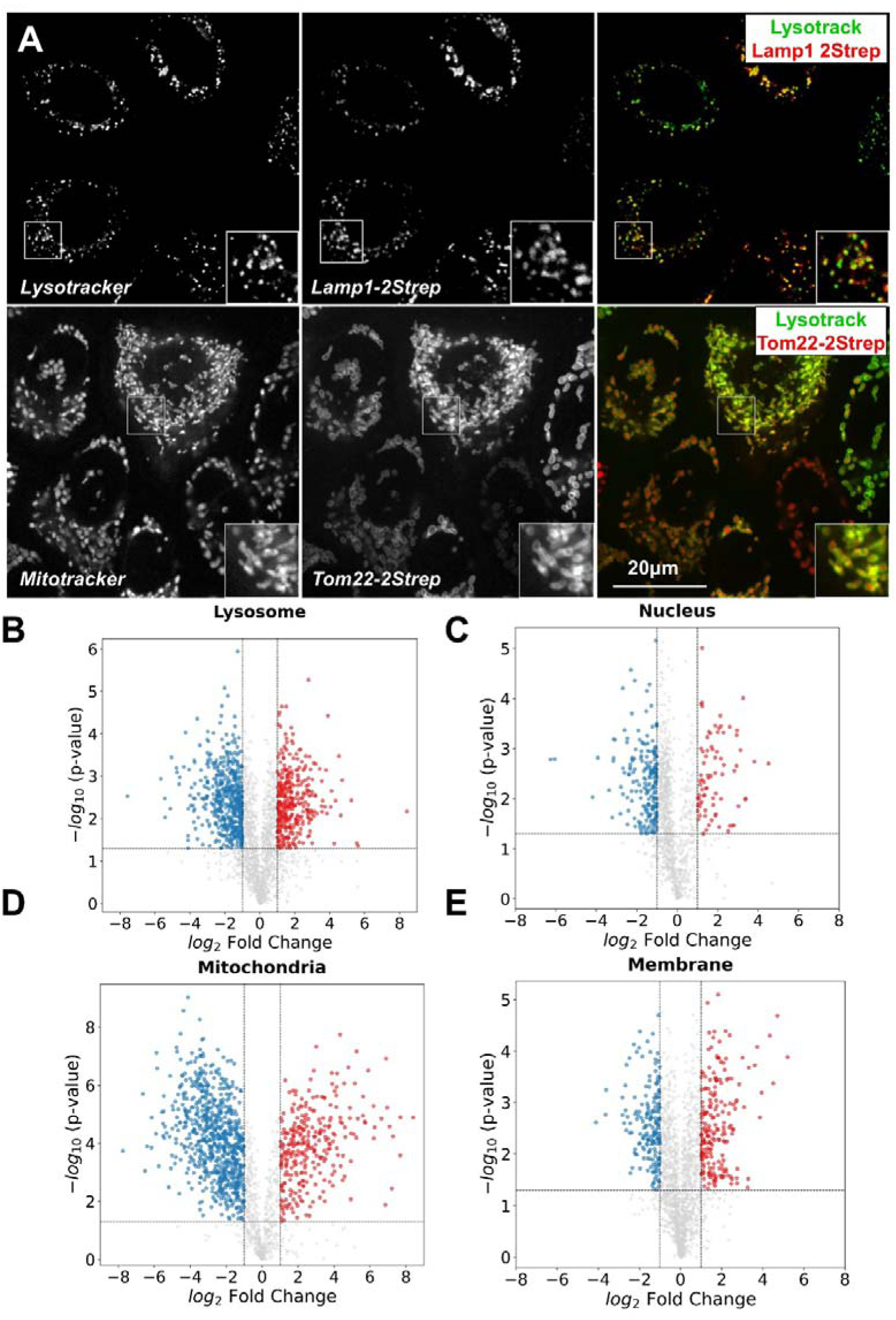
Quantitative Subcellular Proteomics of Keratinocytes. **(A)** Lysosomes and mitochondria were tagged using transgenes encoding integral membrane proteins fused to a Twin-Strep tag: LAMP1 for lysosomes and TOM22 for mitochondria. Live-cell imaging showed strong colocalization of the tagged proteins with LysoTracker and MitoTracker dyes, confirming their correct subcellular localization. **(B–E)** Volcano plots showing proteins enriched in lysosome, nuclear, mitochondria, and plasma membrane enriched fractions from undifferentiated and differentiated keratinocytes.

To validate the functionality of the LC3 with twin-strep tag, keratinocytes expressing tagged LC3 were subjected to nutrient starvation, a condition known to induce autophagosome formation and autophagosome–lysosome fusion. Under these conditions, LC3 puncta exhibited strong colocalization with LysoTracker-positive structures, consistent with enhanced autophagic flux and confirming appropriate localization of the tagged LC3 construct (Supplementary Fig. 2C). In parallel, nuclear and plasma membrane-associated fractions in undifferentiated and differentiated keratinocytes were isolated using established commercial biochemical fractionation approaches.

Quantitative analysis of SILAC-labeled proteomes revealed extensive compartment-specific proteome remodeling during differentiation. In differentiated keratinocytes, we identified 78 nuclear, 208 plasma membrane-associated, 77 LC3-positive vesicle-associated, 357 lysosome-associated, and 293 mitochondrial proteins that were significantly enriched by at least two-fold relative to undifferentiated cells (Supplementary Spreadsheet 1). Our data suggests that epidermal differentiation is accompanied by widespread and coordinated proteomic changes across multiple subcellular compartments.

### Coordinated remodeling of metabolic and vesicular pathways during epidermal differentiation

To identify physiologically meaningful proteome changes associated with keratinocyte differentiation, we analyzed proteins that were significantly enriched based on quantitative SILAC measurements, including proteins exhibiting robust state-specific enrichment despite limited peptide detection in one condition. This approach enabled us to capture both statistically significant changes and highly differential proteins that were reproducibly detected across biological replicates in one cell state but absent or near background in the other.

As an initial validation of this strategy, we examined proteins enriched in undifferentiated keratinocytes within the nuclear fraction. Pathway enrichment analysis revealed strong overrepresentation of proteins involved in cell cycle progression and mitosis (Fig. 3A). Consistently, proteins annotated as mitosis-related exhibited significantly higher abundance in undifferentiated cells compared to differentiated cells (Fig. 3B). These include Cdc23, Cdc40, Cdca7 and Cdk1 which are all well described for their role in regulating the cell cycle. Pcna, which acts as a processivity factor for DNA polymerase delta and is a well-known marker of proliferation, was found to be about 9x more abundant in undifferentiated nuclear fractions. Tp63 is a well characterized regulator of epidermal proliferation and stemness^13,26^. In our nuclear dataset we found significantly higher levels of Tp63 in undifferentiated cells. These findings align with the well-established withdrawal of epidermal keratinocytes from the cell cycle upon differentiation^27^ and indicate that our subcellular proteomics dataset faithfully captures biologically relevant state-dependent changes.

**Figure 3.**
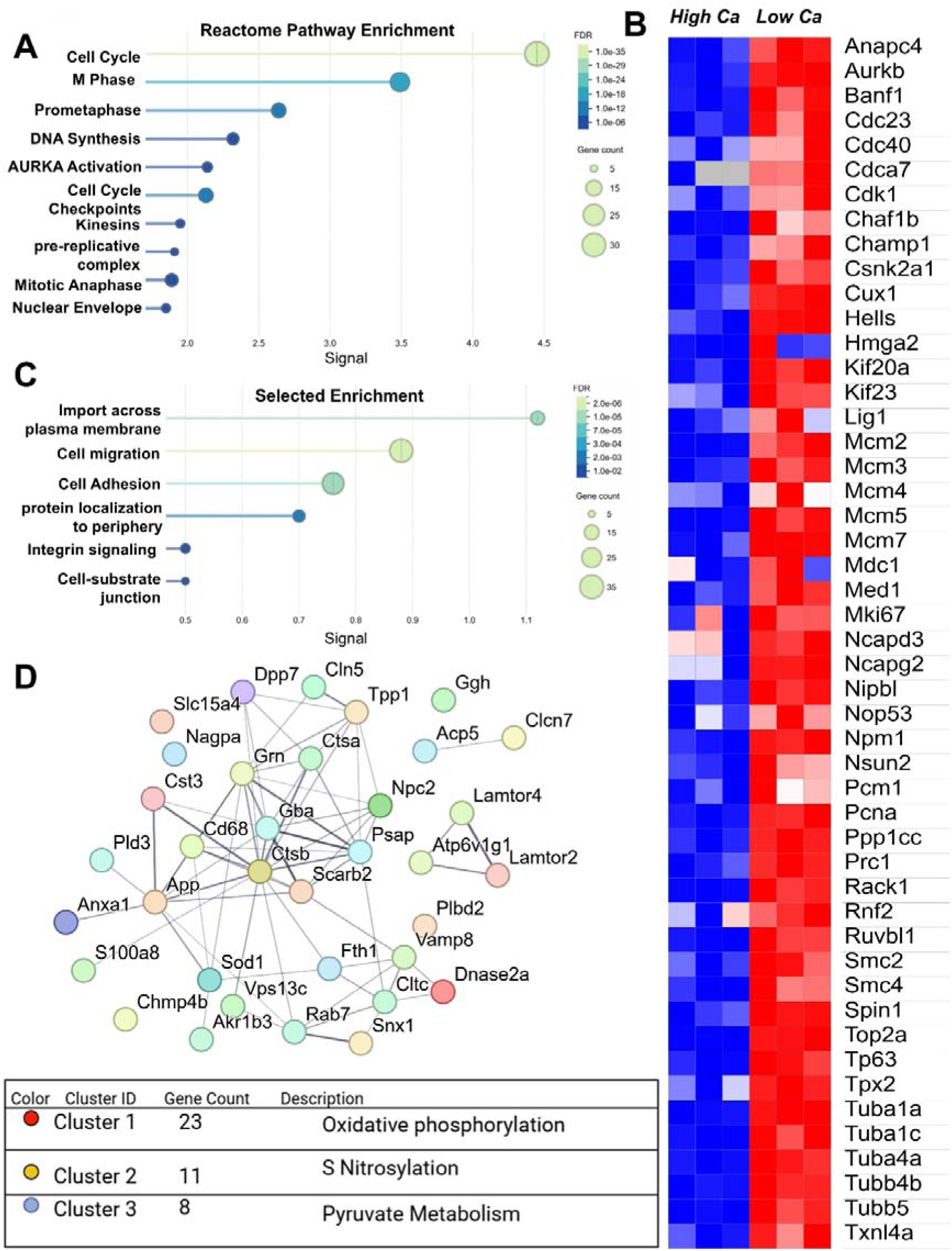
Keratinocyte differentiation is accompanied by coordinated nuclear, membrane, and mitochondrial proteome remodeling. **(A)** Heatmap of proteins from the nuclear dataset which are annotated as cell cycle proteins via string database. A general trend of upregulation in low calcium conditions and downregulation in high calcium conditions is observed. **(B)** Reactome enrichment analysis was performed on proteins identified to be upregulated in undifferentiated cells within the nuclear dataset. **(C)** GO enrichment analysis of membrane proteins identified as upregulated in undifferentiated cells reveals enrichment for terms that are associated with cell-matrix adhesion complexes. **(D)** Cluster analysis on all mitochondria proteins found upregulated in the differentiated state from plasma membrane and mitochondria datasets, along with their respective cluster descriptions, reveals upregulation of proteins controlling oxidative phosphorylation.

Analysis of plasma membrane-associated proteomes demonstrates that proteins enriched in undifferentiated keratinocytes were strongly associated with pathways related to cell migration and integrin-mediated adhesion (Fig. 3C), including components of focal adhesions and hemidesmosomes. This enrichment is consistent with the requirement for proliferative basal keratinocytes to maintain strong adhesion to the underlying basement membrane, a feature that is progressively lost as keratinocytes undergo differentiation/stratification^28–30^.

Given the central roles of mitochondria in cellular metabolism, we analyzed proteome changes in mitochondria during differentiation. To assess mitochondrial remodeling, we analyzed proteins enriched in differentiated cells from both mitochondrial (TOM22–2xStrep) that were annotated as mitochondrial according to the STRING database. We identified 56 annotated mitochondria proteins that were selectively upregulated in differentiated cells in the mitochondria dataset. Clustering analysis revealed that these proteins were predominantly involved in oxidative phosphorylation, pyruvate metabolism, and redox homeostasis (Fig. 3D), suggesting a coordinated shift in mitochondrial metabolic function during keratinocyte differentiation.

Gene ontology analysis of proteins enriched in differentiated cells within the lysosomal fraction revealed strong enrichment for pathways associated with lysosome-dependent catabolic processes and ubiquitin-mediated protein degradation (Fig. 4A). We applied a similar analytical strategy to lysosome-associated proteins enriched in differentiated cells from the LAMP1–2xStrep. 35 proteins annotated as being found at the lysosome were found to be upregulated in differentiated keratinocytes within our lysosome dataset. Unsupervised clustering of these proteins identified three distinct groups (Fig. 4B). One cluster consisted of lysosomal catabolic enzymes, including multiple cathepsin proteases involved in macromolecular degradation. A second cluster was described as trans-golgi network vesicle budding. And the third cluster was associated with mTORC1 regulation. Notably, v-ATPase subunits were consistently increased in differentiated cells across both lysosomal and membrane-associated proteomes (Fig. 4C), suggesting enhanced vesicular acidification capacity during differentiation. Cathepsins are a large class of proteases that are localized to the lysosome and degrade proteins. In our lysosome dataset we saw a general trend of increased cathepsin abundance during differentiation (Fig. 4D). Most cathepsins are activated by the low pH environment so our observations of increased v-ATPase subunits and cathepsin protease abundance suggest that lysosomes are more “active” during keratinocyte differentiation.

**Figure 4.**
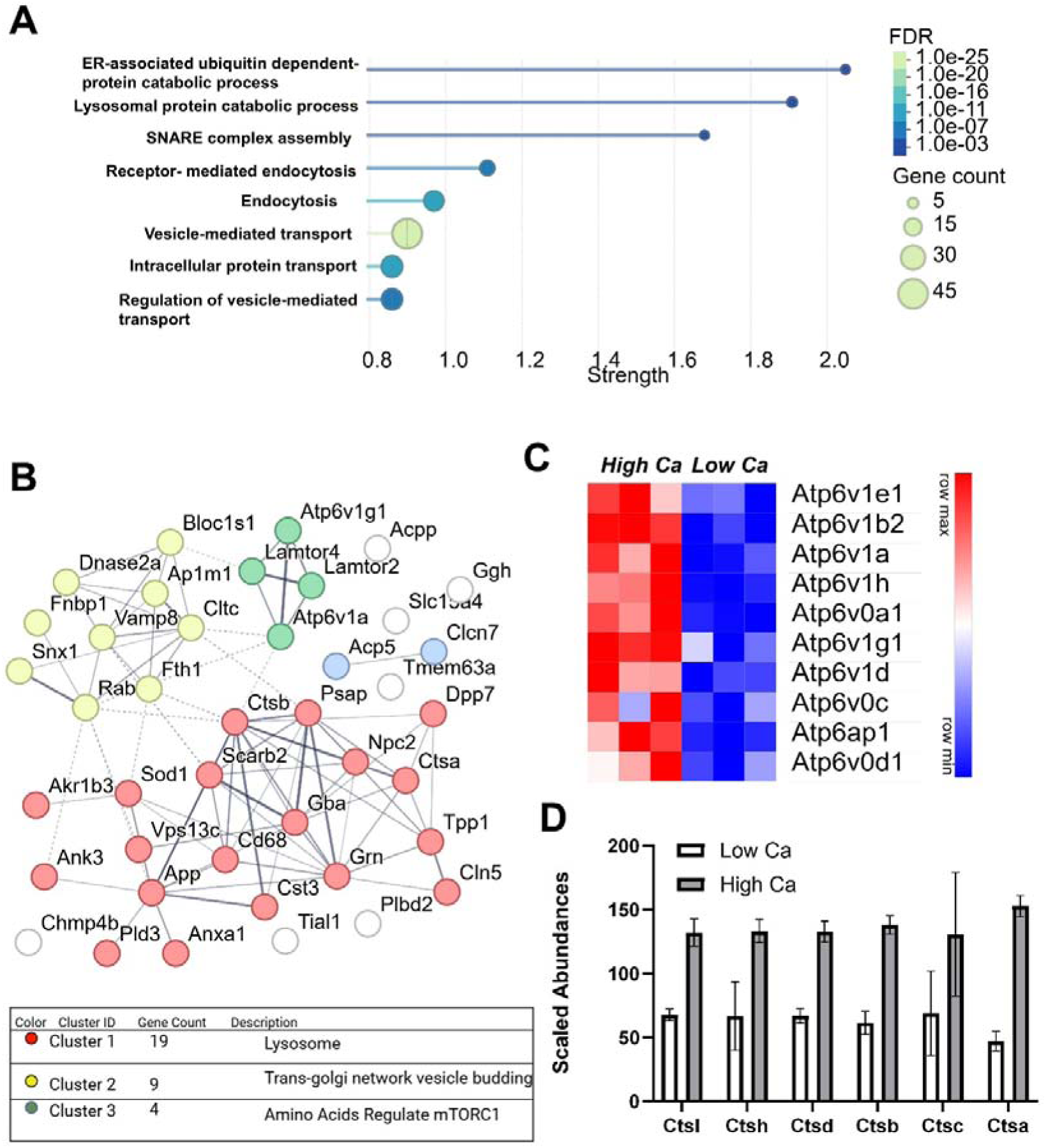
Lysosomal proteomics reveal enhanced lysosome biogenesis and proteolytic activity during keratinocyte differentiation. **(A)** Upregulated proteins in differentiated cells from the lysosome dataset were used for GO analysis. Selected enrichment terms are shown above. **(B)** Proteins found to be upregulated during differentiation in the lysosome dataset were analyzed via the STRING database. K-means clustering was performed on combined entries that were identified as being found or belonging to the lysosomal compartment. **(C)** Heatmap of v-ATPase subunits that were identified in the lysosome dataset shows a general increase in abundance between the two cell states. **(D)** Cathepsin (Cts) proteases and their respective abundances were identified in the lysosome dataset.

Collectively, these quantitative proteomic analyses provide a comprehensive view of compartment-specific proteome remodeling during keratinocyte differentiation and establish a molecular framework for exploring how metabolic and vesicular pathways contribute to epidermal homeostasis.

### Coordinated remodeling of metabolic and vesicular pathways during epidermal differentiation

Our lysosomal proteomics analysis identified many vesicular trafficking proteins that were upregulated in differentiated lysosomal samples (Supplementary Fig. 3A). Vesicular trafficking plays a critical role in regulating receptor signaling and directing cargo to specific intracellular compartments. Notably, several sorting nexin (SNX) proteins were enriched in lysosomes of differentiated keratinocytes (Fig. 5A). SNX proteins comprise a family of endosomal trafficking regulators implicated in receptor sorting and membrane dynamics^31^. Among these, SNX3, a PX domain-only sorting nexin, has been reported to be essential for mammalian development, as *Snx3* knockout results in embryonic lethality in mice^32^. Other SNX proteins have been found to contain redundant functions. *SNX1* and *SNX2* single knockout mice are viable and fertile, but double knockout mice are embryonic lethal indicating redundancy^33^. SNX9 and SNX18 have been shown to contain redundant functions related to clathrin mediated endocytosis^34^. To our knowledge, SNX3 and other sorting nexins have not been previously studied in the context of the epidermis, prompting us to investigate its role in skin keratinocytes.

**Figure 5.**
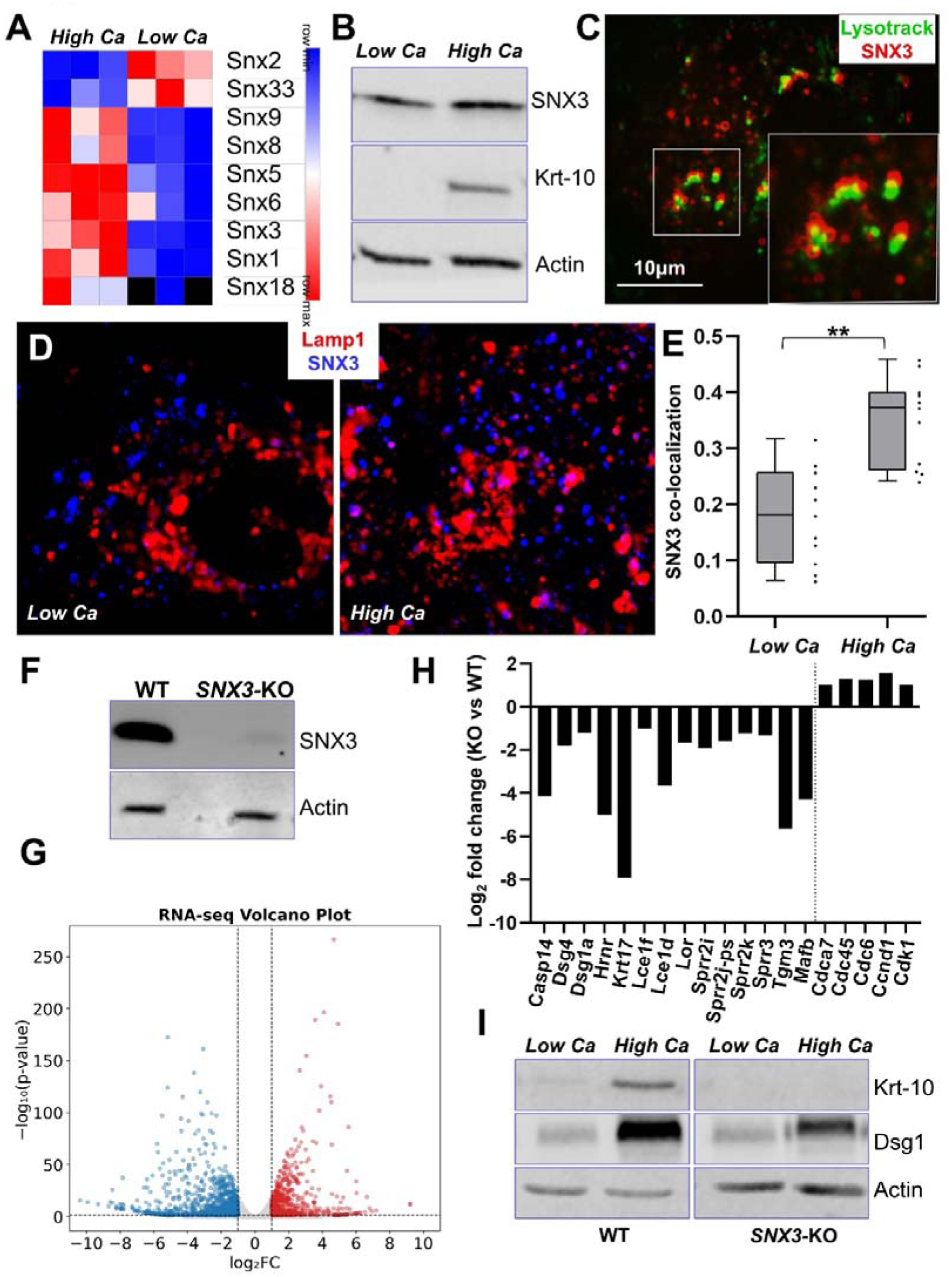
SNX3 regulates Keratinocyte Differentiation. **(A)** Heatmap of all SNX proteins identified in the lysosome dataset. **(B)** SNX3 does not appear to drastically change in total abundance during calcium induced differentiation. **(C)** Cells expressing SNX3-mGFP were stained with lysotracker dye. SNX3 could be found in many instances to be localized to lysotracker labeled vesicles. **(D)** Undifferentiated and differentiated cells were co-stained with Lamp1 and SNX3 antibodies to assess colocalization differences during in vitro differentiation. **(E)** Quantitative analysis of SNX3 signal overlap with Lamp1 signal suggests an increase of SNX3 localization to the lysosome during differentiation. N=11; P<0.05 (Student’s t-test). **(F)** SNX3 gene in mouse keratinocytes was targeted for knockout (KO) using CRISPR-Cas9. Efficient KO was achieved, as determined by western blot. **(G)** Volcano plot of upregulated and downregulated genes in WT and *SNX3* KO cells determined through RNA-seq. **(H)** RNA-seq analysis from *SNX3* KO vs WT cells reveals downregulation of genes associated with differentiation (left of dotted line) and upregulation of cell cycle genes (right of dotted line). All genes shown have an adj p-value<0.05. N=3 biological replicates for both WT and KO cells. **(I)** Western blot analysis of different markers of keratinocyte differentiation upon calcium shift of control and *SNX3* KO cells. Reduction of *Keratin-10* (*Krt-10*) and *Desmoglein 1* (*Dsg1*) expression is observed in *SNX3* KO cells.

The increased representation of SNX3 in lysosomal fractions could reflect either elevated *SNX3* expression or enhanced lysosomal localization during keratinocyte differentiation. However, total SNX3 protein levels remained relatively constant during in vitro differentiation (Fig. 5B). Consistently, skin immunostaining did not reveal a significant increase in SNX3 expression in the suprabasal layers of the epidermis (Supplementary Fig. 3B). SNX3 typically localize at early endosomes^35,36^. To determine whether SNX3 localizes to lysosomes in differentiated keratinocytes, we expressed SNX3-mCherry and LAMP1-mGFP in primary mouse keratinocytes. Interestingly, a substantial fraction of SNX3-containing vesicles were also positive for lysotracker or LAMP1 in differentiated cells (Fig. 5C and Supplementary Fig. 3C). Consistently, colocalization analysis of endogenous SNX3 and LAMP1 demonstrated a significant increase in their spatial overlap during keratinocyte differentiation (Fig. 5D–E). Taken together, these findings indicate that SNX3 is increasingly localized to LAMP1-positive lysosomal vesicles during keratinocyte differentiation.

### Loss of *SNX3* impairs epidermal differentiation

Given that SNX proteins are well-established regulators of protein trafficking, and that SNX3 is enriched in lysosomal fraction upon calcium shift, we hypothesized that SNX3 may contribute to keratinocyte differentiation. To test this, we used CRISPR–Cas9 to generate *SNX3* knockout keratinocytes (Fig. 5F). Bulk RNA-seq analysis of differentiated WT and *SNX3* knockout cells identified 1,064 downregulated and 713 upregulated genes (Fig. 5G and Supplementary Spreadsheet 2). Interestingly, gene ontology analysis revealed that downregulated genes were enriched for differentiation-related processes, whereas upregulated genes were associated with cell cycle pathways (Fig. 5H Supplementary Fig. 4A–B). Deletion of *SNX3* did not appear to significantly change cell morphology or growth of undifferentiated cells *in vitro* (Supplementary Fig. 5A). To directly assess differentiation, we performed in vitro calcium-induced differentiation assays and observed reduced expression of differentiation markers Keratin-10 (Krt-10), Desmoglein 1 (DSG1), and Desmoglein 4 (DSG4) in *SNX3* knockout cells (Fig. 5I and Supplementary Fig. 5B). Together, these data suggest that SNX3 plays an essential role in epidermal differentiation.

### Deletion of *SNX3* promotes cutaneous carcinogenesis *in vivo*

To assess the role of SNX3 *in vivo*, we developed skin organoid cultures with WT or *SNX3* deficient cells and transplanted the skin to nude mice to monitor epidermal differentiation *in vivo.* Surprisingly, while WT grafts healed into normal appearing skin, *SNX3* knockout skin grafts developed abnormal morphology shortly after engraftment, which progressed to ulcerative lesions resembling those observed in cutaneous squamous cell carcinoma (cSCC) (Fig. 6A). Hematoxylin and eosin (H&E) staining of *SNX3*-deficient grafts revealed histopathological features consistent with cSCC, including invasion into the underlying dermis and pronounced parakeratosis (Fig. 6B). In contrast, WT grafts maintained normal epidermal architecture without evidence of tumorigenesis (Fig. 6B).

**Figure 6.**
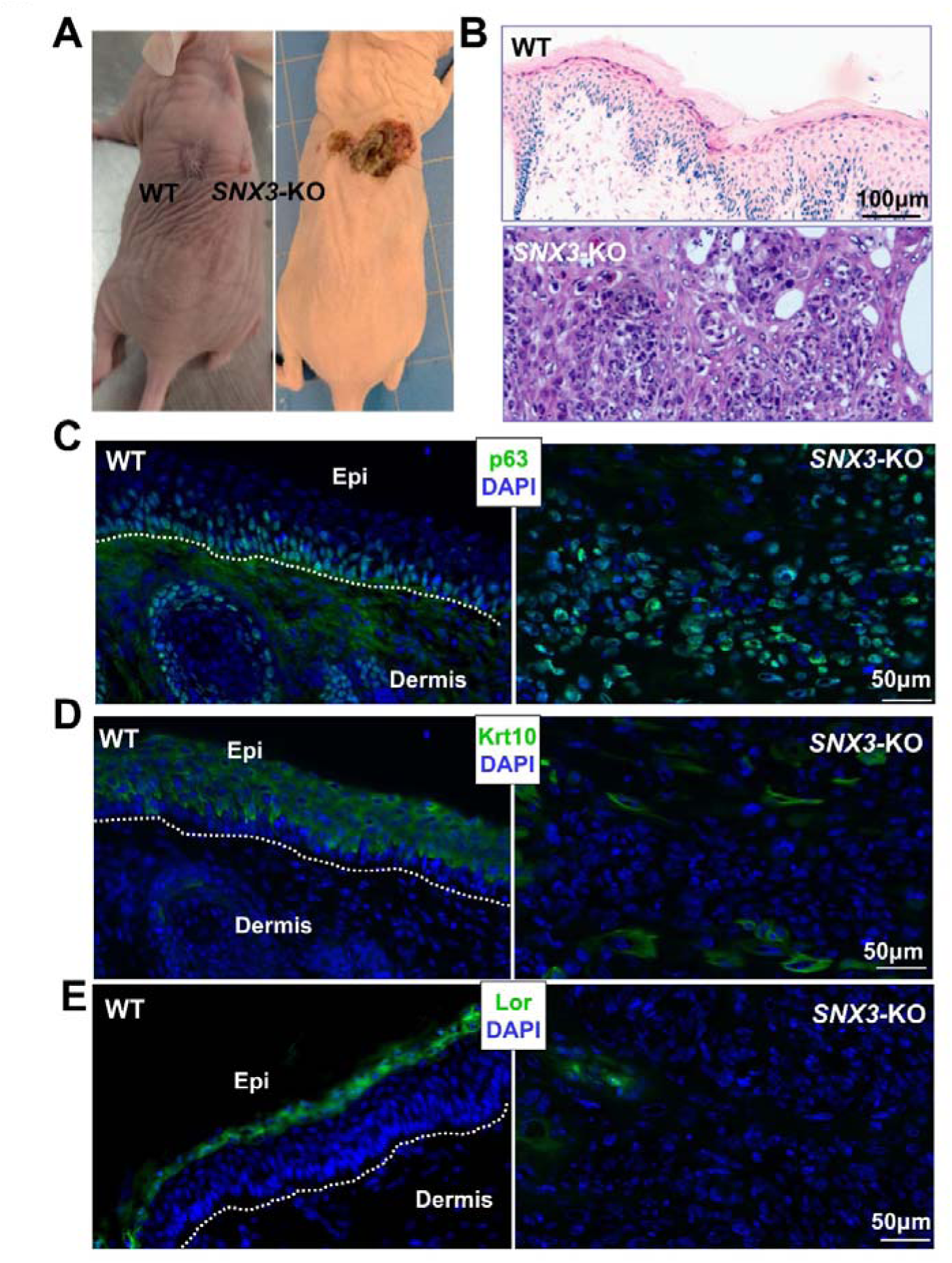
Loss of *SNX3* Leads to Spontaneous Skin Tumorigenesis. **(A)** WT and *SNX3* KO keratinocytes were grafted onto the backs of nude mice. Two weeks post-grafting, WT grafts appeared flat and healed normally, whereas *SNX3* KO grafts displayed an abnormal raised morphology. *SNX3* KO grafts also showed evidence of ulceration and lateral expansion across the mouse back, as shown on the right. **(B)** Histology sections of WT and KO skin grafts. WT grafts exhibit normal epidermal architecture. KO grafts show signs of invasion into the underlying dermis and resemble poorly differentiated cutaneous squamous cell carcinoma. **(C)** P63 levels were assessed via immunofluorescence. In WT grafts P63 is seen predominantly in basal layer cells with expression sharply decreasing in upper layers. In KO grafts, P63+ cells are ubiquitously found in the tumor invading into the dermis. **(D)** Expression of *Krt-10* in WT and KO grafts assessed via immunofluorescence shows a reduction in cells expressing *Krt-10* upon loss of *SNX3*. **(E)** Expression of *loricrin in* WT and KO grafts assessed via immunofluorescence shows a lack of *loricrin* expression in KO tumors.

p63-positive cells are typically restricted to the basal layer of the epidermis and decline markedly as cells undergo suprabasal differentiation^26,37,38^. Immunostaining revealed a substantial expansion of p63⁺ cells within the invasive regions of *SNX3* knockout grafts, consistent with poorly differentiated cSCC phenotypes^39^ (Fig. 6C). Furthermore, Krt-10-expressing cells were reduced in *SNX3* knockout skin, and loricrin staining was nearly absent, indicating impaired terminal differentiation (Fig. 6D–E). Immunofluorescence staining for Ki67 showed proliferation confined to the basal layer in WT grafts, whereas Ki67⁺ cells were broadly distributed throughout the tumor mass in *SNX3* knockout grafts (Supplementary Fig. 5C). Taken together, these data indicate that loss of SNX3 in keratinocytes inhibits epidermal differentiation and drives invasive, cSCC-like tumorigenesis *in vivo*.

### SNX3 regulates Notch signaling in epidermal keratinocytes

Notch signaling is a well-established driver of keratinocyte differentiation and functions as a tumor suppressor in skin^12^. Notch signaling output depends on proteolytic cleavage of Notch receptors to generate the transcriptionally active NICD^40^. Importantly, trafficking of Notch receptors through the endosomal system is a key regulator of downstream signaling activity^41,42^. Given that SNX proteins are established regulators of receptor trafficking, we hypothesized that SNX3 may modulate Notch signaling.

To test this hypothesis, we transduced keratinocytes with a Notch reporter in which luciferase expression is driven by CBF1/RBP-Jκ response elements^43^. *SNX3* knockout cells exhibited significantly reduced luciferase activity compared to WT controls (Fig. 7A). Although multiple members of the Notch family are expressed in the skin, Notch1 appears to be particularly important. Genetic ablation of Notch1 results in significant phenotypic abnormalities such as impaired differentiation and tumorigenesis whereas deletion of the other Notch receptors has no or minimal phenotype^14,44^. Co-immunostaining of Notch1 and SNX3 revealed discrete Notch1-positive puncta associated with SNX3 in differentiating keratinocytes (Fig. 7B).

**Figure 7.**
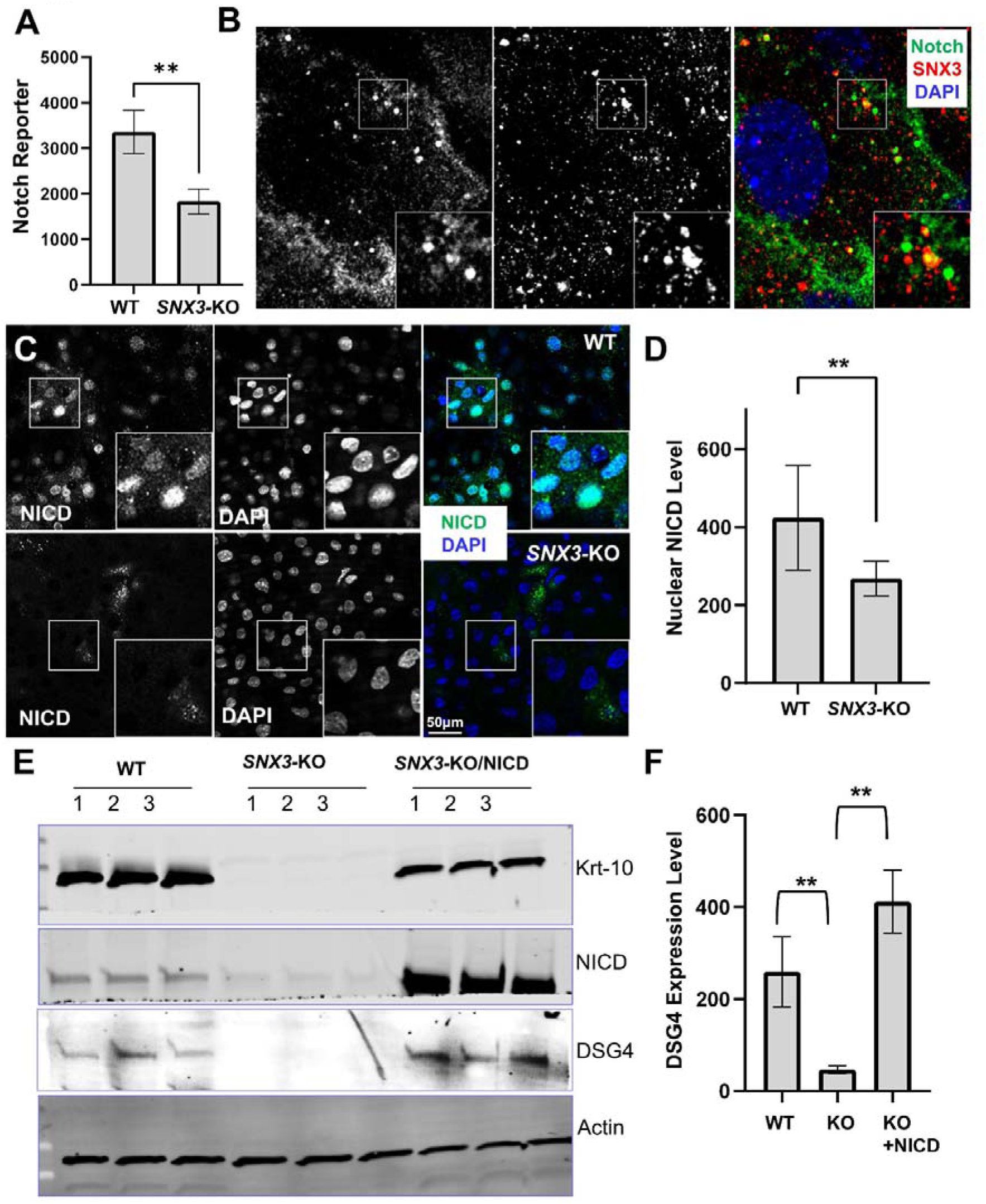
SNX3 Regulates Notch Signaling During Epidermal Differentiation. **(A)** WT and *SNX3* KO cells were transduced with a notch reporter driving luciferase expression. Lower reporter activity was observed in the *SNX3* KO cells upon calcium shift. N=3 biological replicates; P<0.05 (Student’s t-test). **(B)** Keratinocytes were co-stained with Notch1 and SNX3 antibodies. Notch1 intracellular puncta co-localizes with SNX3. **(C)** Immunofluorescence staining of NICD suggests lower levels of NICD in the nucleus of SNX3 KO cells. **(D)** Immunofluorescence quantitative analysis of nuclear Notch1 intensity from in WT and *SNX3* KO cells. N=50, and p<0.05 (Student’s t-test). **(D)** Notch signaling was rescued by ectopic expression of NICD in *SNX3* KO cells. Differentiation markers Krt-10 and DSG4 expression was assessed via western blot. **(E)** DSG4 protein levels were quantified via western blot densitometry. N=3, and ** p<0.05 (Student’s t-test).

Notch receptor cleavage and signaling are influenced by endolysosomal trafficking and vesicular acidification^45,46^. Prior studies have shown that impaired endolysosomal acidification reduces Notch1 cleavage and signaling output^45–49^. To assess whether SNX3 loss affects this process, we measured lysosomal pH using a pH-sensitive dye (Supplementary Fig. 6A). Mean fluorescence intensity was similar between WT and *SNX3* knockout cells, indicating that lysosomal acidification is not significantly altered (Supplementary Fig. 6B). In addition, the cellular distribution of full-length Notch1 and its ligand Delta-like canonical Notch ligand 1 (DLL1) was not noticeably altered between WT and knockout cells (Supplementary Fig. 6C–D), suggesting that trafficking of the full-length receptor and ligand is normal. By contrast, immunostaining for cleaved NICD followed by quantification of nuclear signal level revealed markedly reduced nuclear NICD in *SNX3*-deficient cells (Fig. 7C-D). Ectopic expression of NICD in *SNX3* knockout cells significantly rescued the expression of multiple differentiation markers, suggesting that reduced Notch cleavage and NICD-dependent nuclear signaling contribute to the role of SNX3 in epidermal differentiation (Fig. 7E-F).

Together, these findings suggest that SNX3 regulates Notch signaling not through global changes in lysosomal acidification, but likely by facilitating proper trafficking of the Notch1 receptor through the endolysosomal system (Supplementary Fig. 7).

## DISCUSSION

Epidermal differentiation is a highly coordinated process in which proliferative basal keratinocytes permanently exit the cell cycle and progressively acquire the specialized structural and biochemical properties required for barrier formation^2,9,10^. While transcriptional programs governing this transition have been extensively studied, less is known about how intracellular organelles are remodeled to support differentiation-associated changes in signaling, metabolism, and cellular architecture. In this study, we used compartment-resolved quantitative proteomics to define organelle-specific proteome remodeling during keratinocyte differentiation. Our analyses reveal widespread and coordinated changes across lysosomes, mitochondria, autophagic vesicles, plasma membrane, and nucleus, indicating that epidermal differentiation is accompanied by global reorganization of subcellular systems rather than by isolated changes in a few pathways. Importantly, this organelle-centered approach led to the identification of SNX3 as a previously unrecognized regulator of epidermal differentiation and skin tissue homeostasis.

Our data suggest that differentiation is associated with a broad shift in both metabolic state and vesicular function. Mitochondrial proteomics revealed enrichment of proteins linked to oxidative phosphorylation, pyruvate metabolism, and redox regulation, consistent with the idea that differentiating keratinocytes undergo metabolic rewiring rather than simply shutting down biosynthetic activity^50–52^. In parallel, lysosomal fractions from differentiated cells were enriched for v-ATPase subunits, cathepsins, and proteins involved in vesicle trafficking and lysosome-associated catabolism. The v-ATPase is a multi-subunit complex consisting of two domains V_0_ and V_1_ which must assemble together to achieve lysosomal acidification^53,54^. These observations, together with the increased cathepsin activity detected in differentiated keratinocytes, support a model in which lysosomal activity is enhanced during differentiation. Such changes are likely to be important for the turnover of macromolecules and organelles as keratinocytes progress toward terminal differentiation. More broadly, these results highlight the value of subcellular proteomics in identifying regulatory processes that are not readily captured by transcriptomic analyses alone, particularly in contexts where protein localization and organelle association are likely to be functionally important.

Among the differentiation-associated changes identified in our lysosomal dataset, SNX3 emerged as a particularly compelling candidate. Sorting nexins are well known regulators of membrane trafficking, receptor sorting, and endosomal dynamics, yet their functions in epidermal biology remain poorly understood. Mechanistically, our data support a model in which SNX3 promotes epidermal differentiation at least in part through regulation of Notch signaling. Notch is a central driver of keratinocyte differentiation and functions as a tumor suppressor in the epidermis^14,55–57^. *SNX3* loss reduced Notch reporter activity and diminished nuclear NICD accumulation. Moreover, ectopic NICD expression rescued the differentiation defect in *SNX3*-deficient cells, indicating that impaired Notch activation contributes substantially to the phenotype. Notably, we did not observe major changes in overall lysosomal acidification or alterations in the steady-state distribution of full-length Notch1 or DLL1, arguing against a model in which SNX3 broadly disrupts lysosome function or receptor expression. Notch receptor signaling is tightly coupled to membrane trafficking, proteolytic processing, and vesicular maturation, and perturbations in endolysosomal organization can strongly influence downstream signaling output. Our results extend this concept to epidermal differentiation by identifying SNX3 as a trafficking factor that connects differentiation-associated organelle remodeling with Notch-dependent keratinocyte differentiation. More broadly, these findings raise the possibility that dynamic reorganization of organelles is not merely a consequence of differentiation, but an active component of the differentiation program itself. In this view, remodeling of lysosomes and related vesicular compartments could create a permissive intracellular environment for the execution of Notch signaling. In support of this idea, we consistently saw an increase in the abundance of v-ATPase subunits within our differentiated lysosome samples. Proper lysosomal acidification has been suggested to be necessary for Notch signaling in both mouse and fly^36,37,39,40,46^. It is unclear how regulation of v-ATPase endolysosome assembly during keratinocyte differentiation contributes to Notch signaling. Future studies will be essential to uncover the link between v-ATPase dynamics and Notch signaling, particularly regarding their collective impact on skin biology and disease. Notably, it is important to mention that knockdown of V-ATPase subunits or pharmacological inhibition of the V-ATPase has been shown to cause severe disorganization and differentiation defects in human skin organoids^8^.

An additional implication of our findings is the link between defective organelle-associated trafficking and tumorigenesis in stratified epithelium. The spontaneous development of invasive cSCC-like lesions in *SNX3*-deficient grafts underscores the importance of proper differentiation for suppressing malignant transformation in the skin. Given the well-established role of Notch signaling in promoting keratinocyte differentiation and suppressing skin tumorigenesis^14,55–57^, reduced Notch activity likely contributes to the phenotype of *SNX3*-deficient epidermis. However, the severe *in vivo* abnormalities observed upon *SNX3* loss also suggest that additional trafficking-dependent pathways may be involved. Future studies will be needed to define the molecular machinery downstream of SNX3 and to determine how its dysfunction contributes to epidermal differentiation defects and cutaneous carcinogenesis.

In summary, our study identifies SNX3 as a regulator of epidermal homeostasis and raises the possibility that differentiation-associated organelle remodeling contributes directly to Notch signaling control. These findings provide a framework for further dissecting how intracellular trafficking pathways shape epidermal cell fate.

## STAR METHODS

### KEY RESOURCES TABLE

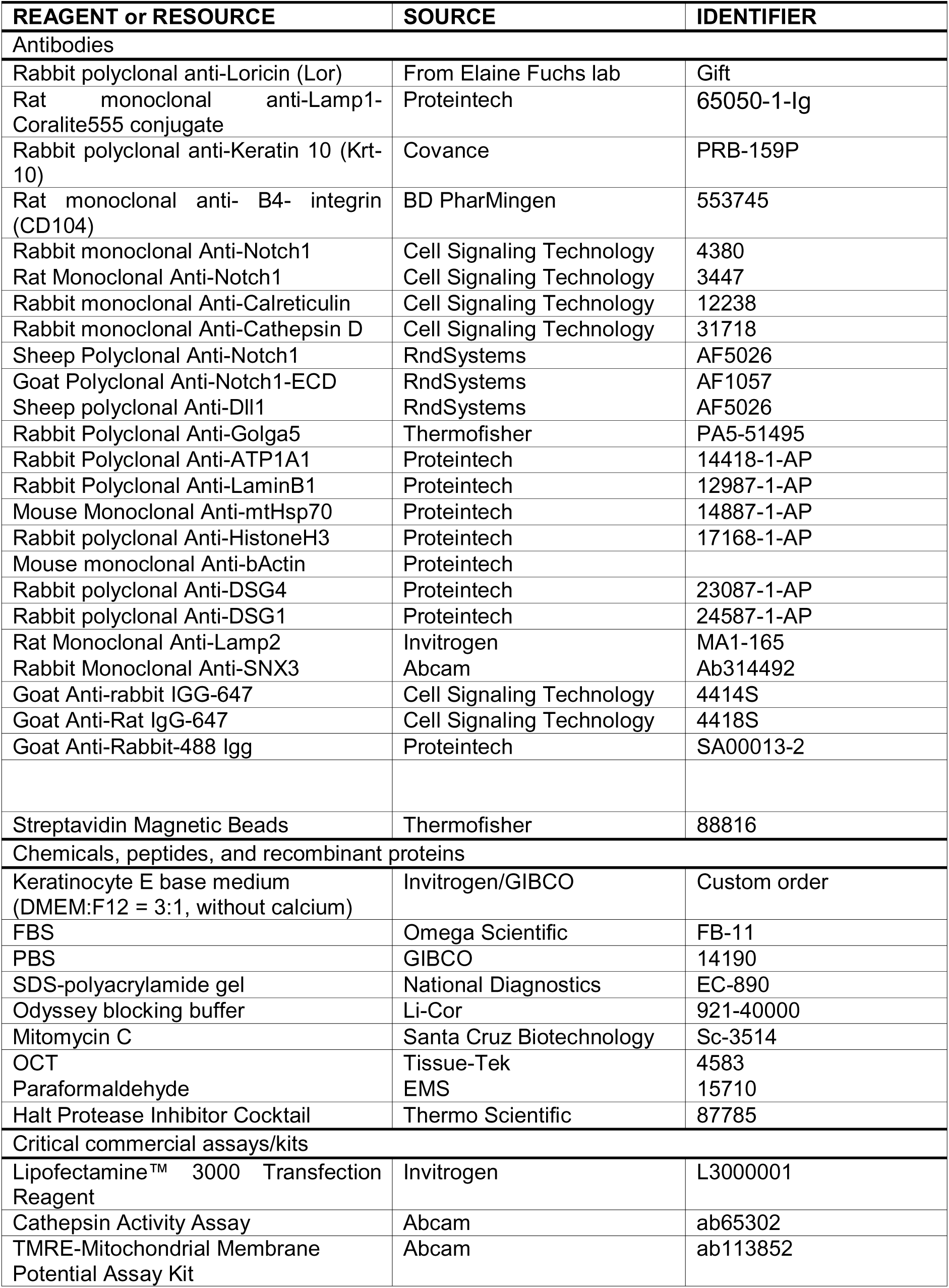
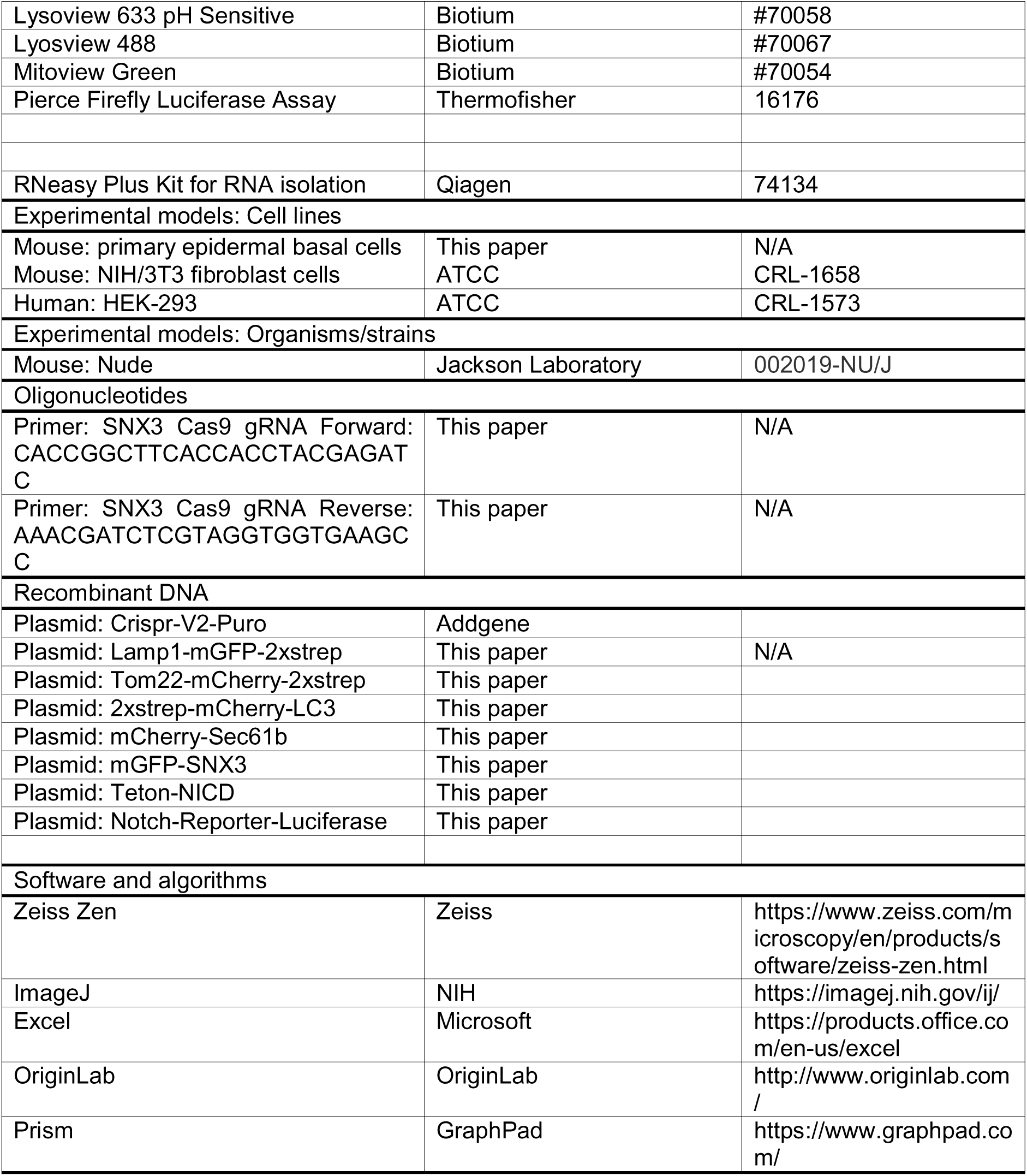

### RESOURCE AVAILABILITY

#### Lead contact and Materials availability

Further information and requests for resources and reagents should be directed to and will be fulfilled by the Lead Contact, Xiaoyang Wu.

### EXPERIMENTAL MODEL AND STUDY PARTICIPANT DETAILS

#### Animals

Athymic nude mice (strain # 002019) from Jackson Laboratories were used for skin grafting experiments. CD1 mice were used to obtain primary mouse keratinocytes. Around 8-week-old female nude mice were used for skin grafting. All the mice were housed under pathogen-free conditions in ARC (animal resource center) of University of Chicago under a 12 hr light-dark cycle. All the subjects were not involved in any previous procedures. The experimental protocol was reviewed and approved by the Institutional Animal Care and Use Committee of University of Chicago.

#### Primary Epidermal Cell Culture

Primary mouse keratinocytes were isolated using previously reported methods (Lee et al., 2017). Epidermis of newborn mice or E18.5 was separated from dermis by an overnight treatment with dispase. Then, the primary keratinocytes were dispersed from the epidermis using trypsin. Keratinocytes were co-cultured with mitomycin C-treated 3T3 fibroblast feeder cells until the third passage. Cells were maintained in E-media supplemented with 15% FBS. The final concentration of Ca^2+^ is 0.05 mM. High calcium shift was performed using E-media supplemented with 15% FBS, with Ca^2+^ at a final concentration of 1.5 mM. HEK293 cells were cultured in DMEM supplemented with 10% FBS.

### METHOD DETAILS

#### Plasmid DNA Constructions

Cas9-SNX3 lentiviral vector was made by annealing SNX3gRNA-forward (CACCGGCTTCACCACCTACGAGATC) and SNX3gRNA- reverse (AAACGATCTCGTAGGTGGTGAAGCC) oligos and inserting the annealed product into lenti-CrisprV2-Puro. Coding sequences for Lamp1, Tom22, Sec61b, and SNX3 were obtained from Horizon discovery and then subcloned into our in-house lentiviral vector.

#### Skin organotypic culture and grafting

Skin organotypic culture and grafting were performed as previously described^58^. Decellularized dermis (1 × 1 cm square shape) was prepared from newborn CD1 mice skin via EDTA treatment^59^. 2 × 106 cultured keratinocytes with desired genomic modifications were seeded onto the dermis in cell culture insert. Then, the skin culture was exposed to air/liquid interphase after an overnight attachment to form skin organoids. For grafting with skin organoids, nude mice aged ∼12 weeks were anaesthetized. Two 1 × 1 cm square shape wounds were introduced to the back skin of the nude mice. After transplantation of skin organoids to the fresh wounds, the wound edge was sealed with surgical glue. The animals with skin grafts were housed separately, and the bandages over the wound could be removed 1 week after surgery^58,60,61^. All the experiments were repeated more than three times (three biological replicates). For phenotypic analysis using immunostaining, at least three sections were taken from each graft for quantifications.

#### Protein biochemical analysis

Western blot was conducted as previously described^62^. Cell lysates were prepared with RIPA buffer containing protease inhibitors. After the concentration of total protein is assessed, equal amounts of the cell lysates were resolved in sodium dodecyl sulfate–polyacrylamide gel and electroblotted onto a nitrocellulose membrane. The membrane was incubated with Odyssey blocking buffer (Li-COR biosciences, Lincoln, NE) for 1 h at room temperature, followed by an overnight incubation with desired primary antibody at 4°C. Immunoblots were washed three times with 1× Tween 20/phosphate-buffered saline (PBST) and incubated with secondary antibody (1:10,000 dilution) at room temperature for 1 h. Blot was washed with 1× PBST for another three times. LI-COR Odyssey scanner was used to visualize the blotting signals, and LI-COR Biosciences Software was used to conduct the quantification.

#### Histology and immunofluorescence

Skin or wound samples were embedded in OCT, sectioned, and fixed in 4% paraformaldehyde. Antibodies were diluted according to manufacturer’s instruction unless indicated.

Images were collected on a Zeiss AxioObserver7 and analyzed using Zeiss Zen software and ImageJ. Live cell imaging and cellular immunofluorescence images were collected using a SoRa Subdiffraction Marianas Spinning Disc Confocal. Colocalization analysis was performed using the FIJI plugin JaCoP.

#### Notch Reporter Assay

The Pierce firefly luciferase assay kit was used for notch reporter activity assessment. WT and KO cells expressing the Notch reporter were lysed in provided cell lysis buffer. Protein concentration was equilibrated based on BCA measurements. Equal amounts of protein from both samples were used for measuring bioluminescence.

#### Mitochondria TMRE Assay

30000 keratinocytes were plated in the wells of a black 96 well plate. One group was then differentiated by switching the medium to high calcium for 24 hours. TMRE fluorescence was measured using a Synergy Neo HST Plate Reader. To control for differences in cell numbers that could occur as the undifferentiated cells continue dividing, we used a nuclear stain as a normalization factor.

#### Cathepsin Activity Assay

Cell lysates were collected from undifferentiated diving progenitors and differentiated keratinocytes. The abcam cathepsin activity assay kit was used to detect the level of cathepsin activity in the samples. Equal protein amounts (∼20ug) were used for the assay. Fluorescence measurements were obtained using a Synergy NEO HST Plate Reader.

#### Subcellular fractionation and Organelle Immunoprecipitation

Membrane enrichment was performed using the Invent Biotechnologies membrane and cell fractionation kit according to the manufacturer’s instructions. Nuclear enrichment was performed using the Abcam cell fractionation kit according to the manufacturer’s instructions. Organelle immunoprecipitation was performed by homogenizing cells in PBS with a dounce homogenizer. Cell extracts were then centrifuged at 1000xg for 5 minutes. The supernatant was collected and the centrifugation step was repeated once more. The supernatant was then mixed with 100uL of streptavidin magnetic beads and incubated on a tube rotator for 15 minutes. Following incubation, the beads were washed three times with ice cold PBS. Elution was performed by incubating the beads in 1%SDS and heating at 95°C for 5 minutes.

#### SILAC Sample Preparation

Cells were cultured in heavy (K8+R6) or light (K0+R0) SILAC media for 10 doublings to fully label proteomes prior to collection. Organelles were fractionated as described above. Protein concentrations from heavy and light organelle fractions were quantified using the Bradford protein assay (Bio-Rad Laboratories, Hercules, CA). Equal amounts (1:1, w/w) of heavy and light proteins were mixed and digested with trypsin (1:50, Promega, Madison, WI) for 16.5 h in 50 mM ammonium bicarbonate (Sigma-Aldrich, St. Louis, MO). The resulting tryptic peptide mixtures were directly subjected to HPLC–MS/MS analysis.

#### HPLC–MS/MS Analysis

Peptides were loaded onto a homemade capillary C18 column (10 cm length × 75 mm ID, 3 µm particle size, Maisch GmbH, Beim Bruckle, Germany). HPLC–MS/MS analysis was performed on an Orbitrap Exploris 480 mass spectrometer (Thermo Fisher Scientific, Waltham, MA) coupled to an EASY-nLC 1000 system (Thermo Fisher Scientific). Mobile phase A consisted of 0.1% formic acid in water, and mobile phase B was 0.1% formic acid in acetonitrile. Peptides were separated with a linear gradient from 2% to 35% B over 90 min at a flow rate of 0.3 μL/min.

Mass spectra were acquired in positive-ion mode across the m/z range 300–1,100 at a resolution of 60,000. Data-dependent MS/MS was performed by selecting the 20 most intense precursor ions for higher-energy collisional dissociation (HCD, NCE 30) at a resolution of 15,000. Acquisition parameters were as follows: isolation window, 1.6 m/z; default charge state, 2+; normalized collision energy, 30%; maximum injection time, auto; first mass, 100 m/z; AGC target, standard; dynamic exclusion, 20 s (after two occurrences).

#### Data Analysis

Raw files were processed with Proteome Discoverer 2.5.0.400 (Thermo Fisher Scientific) and searched against the UniProt mouse reference FASTA database (downloaded March 2021) supplemented with common contaminants using the Sequest HT search engine. Static carbamidomethylation of cysteine (+57.021 Da) was set as a fixed modification, while oxidation of methionine (+15.995 Da) and acetylation of protein N-termini (+42.011 Da) were included as variable modifications. Trypsin was specified as the digestion enzyme, allowing up to two missed cleavages. Precursor and fragment mass tolerances were set to 10 ppm and 0.02 Da, respectively.

For SILAC quantification, a static heavy label modification of +8.014 Da on lysine was specified. Peptide-spectrum matches were validated with Percolator, and results were filtered to a false discovery rate (FDR) < 1% at both peptide and protein levels, with a requirement of at least one unique peptides per protein. Quantification of heavy-to-light ratios was performed using the Precursor Ions Quantifier node, with MS1-based area integration and normalization by the total peptide amount.

### QUANTIFICATION AND STATISTICAL ANALYSIS

Statistical analysis was performed using Excel or GraphPad Prism software. Box plots are used to describe the entire population without assumptions on the statistical distribution. Student’s *t*-test was used to assess the statistical significance (*P*-value) of the difference for most experiments. *p* < 0.05 was considered statistically significant. *p* < 0.05 is indicated as one asterisk (*), and *p* < 0.01 is indicated as two asterisks (**).

#### RNA Seq Analysis

RNA was isolated from WT and knockout cells using the Qiagen RNeasy Plus Kit. Bulk RNA sequencing was performed by Novogene. QC was performed by removing reads containing adapters, removing reads containing N>10% (N represents bases that could not be determined), and removing low quality reads. Mapping was performed using HISAT2 aligning reads to mouse transcriptome. Differential expression analysis was performed using DESEQ2. Upregulated and downregulated genes were used for gene ontology analysis. RNA-seq volcano plots were made using python.

#### Proteomics Analysis

Upregulated and downregulated proteins were selected if they had a LogFC>1 or <-1 and p-value<0.05. For proteins that did not have p-values due to missing data in one of the trials, or proteins not found in one condition we selected these proteins based on two criteria: the protein had to be detected 3 times in the light or heavy conditions and logFC>1 or <-1. Differentially abundant proteins were uploaded to the string database for enrichment analysis, network modeling, and cluster analysis.

## DATA AND SOFTWARE AVAILABILITY

A complete list of software for data analysis and processing can be found in the Key Resources Table.

## Acknowledgments

We thank Dr. Elaine Fuchs at the Rockefeller University for sharing reagents. We thank Linda Degenstein from the University of Chicago transgenic core facility for her excellent technical assistance. The animal studies were carried out in the ALAAC-accredited animal research facility at the University of Chicago. This work was supported by NIH grants R01OD023700, R21AR080761, R01DA047785, R01AR78555, R34AR084090, Leo foundation research grant (to XW) and NIH R01AR78555 (to YZ).

## Data Availability Statement

For this study, we will make our data available to the scientific community, which will avoid unintentional duplication of research. All the research data will be shared openly and in a timely manner in accordance with the most recent NIH guidelines (http://grants.nih.gov/grants/policy/data_sharing/).

## Author Contributions

XW, AH, and YZ designed the experiments. AH, XS, JG, JL, HL, BT, and SS performed the experiments. AH, XS, and XW analyzed the data. AH and XW wrote the manuscript. All authors edited the manuscript.

