## Supplementary-Figs for "A proteomic atlas of organelle remodeling identifies lysosomal SNX3 as a regulator of Notch signaling in epidermal differentiation"

**Supplementary Information**

**Supplementary Figure 1. Inhibition of Lysosomal Acidification Inhibits Keratinocyte Differentiation. (A)** Keratinocytes were treated with bafilomycin A1 to inhibit lysosomal acidification. Western blotting was performed to assess the effect of lysosomal inhibition on the keratinocyte. **(B)** Quantitative analysis showed a significant decrease in loricrin protein levels following bafilomycin A1 treatment. n = 4; ** P < 0.05, Student’s t-test.

**Supplementary Figure 2. Isolation of subcellular compartments using Twin-Strep–tagged marker proteins. (A–B)** Lysosomes and mitochondria were isolated from mouse primary keratinocytes expressing LAMP1–2×Strep or TOM22–2×Strep using streptavidin magnetic beads. Compartment-enriched samples were evaluated by western blotting with the indicated antibodies. **(C)** Keratinocytes expressing LC3 fused to mCherry and 2×Strep tags were starved and stained with LysoTracker. The mCherry-LC3 signal strongly overlapped with the LysoTracker signal, verifying the functionality of the LC3 construct.

**Supplementary Figure 3. SNX3 is enriched in vesicular compartments during keratinocyte differentiation. (A)** STRING network analysis of vesicular proteins encoded by genes identified as upregulated in differentiated cells in the lysosome dataset. **(B)** Immunostaining of SNX3 in neonatal mouse skin. β4 integrin marks the basement membrane. **(C)** Live-cell imaging of keratinocytes expressing SNX3-mCherry and LAMP1-mGFP.

**Supplementary Figure 4. Enrichment Analysis from RNA-Seq Data of WT vs *SNX3* KO Cells. (A)** Reactome enrichment analysis of downregulated genes in *SNX3* KO cells reveals enrichment for terms related to epidermal differentiation. **(B)** Reactome enrichment analysis of upregulated genes in *SNX3* KO cells reveals enrichment for genes related to cell proliferation.

**Supplementary Figure 5. SNX3 regulates keratinocyte differentiation. (A)** Morphology of WT and *SNX3* KO keratinocytes *in vitro*. **(B)** Desmoglein 4 (DSG4) immunoblot of WT and *SNX3* KO cells after calcium shift. DSG4 levels are strongly reduced. N=3; P<0.05 (Student’s t-test). **(C)** WT and *SNX3* KO skin grafts were immunostained for Ki67. In WT grafts, Ki67-positive cells were largely confined to the basal layer, whereas in *SNX3* KO tumors, Ki67-positive cells were distributed throughout the tumor mass.

**Supplementary Figure 6. *SNX3* KO cells retain normal lysosomal pH and Notch1/DLL1 localization.** **(A-B)** Lysosomal pH were assessed using the pH sensitive fluorescent dye lysoview (quantification in B). Lysosomal acidification is not significantly altered in *SNX3* KO cells. **(C)** Immunostaining of Delta-like canonical Notch ligand 1 (Dll1), the Notch ligand, in WT and *SNX3* KO cells. Expression and localization of Dll1 appears to be similar between WT and KO cells. **(D)** Immunostaining of Notch1 using an antibody that targets the extracellular domain of the protein. Expression and cell surface localization of Notch1 appears to be similar between WT and KO cells.

**Supplementary Figure 7. A working model in which SNX3 promotes Notch signaling**. SNX3 functions at the early endosome to facilitate Notch1 trafficking to the endolysosomal compartment, where Notch1 is cleaved by γ-secretase to release transcriptionally active NICD. Nuclear NICD activates downstream Notch target genes, thereby promoting keratinocyte differentiation, restricting inappropriate basal-like proliferation, and supporting normal epidermal stratification.

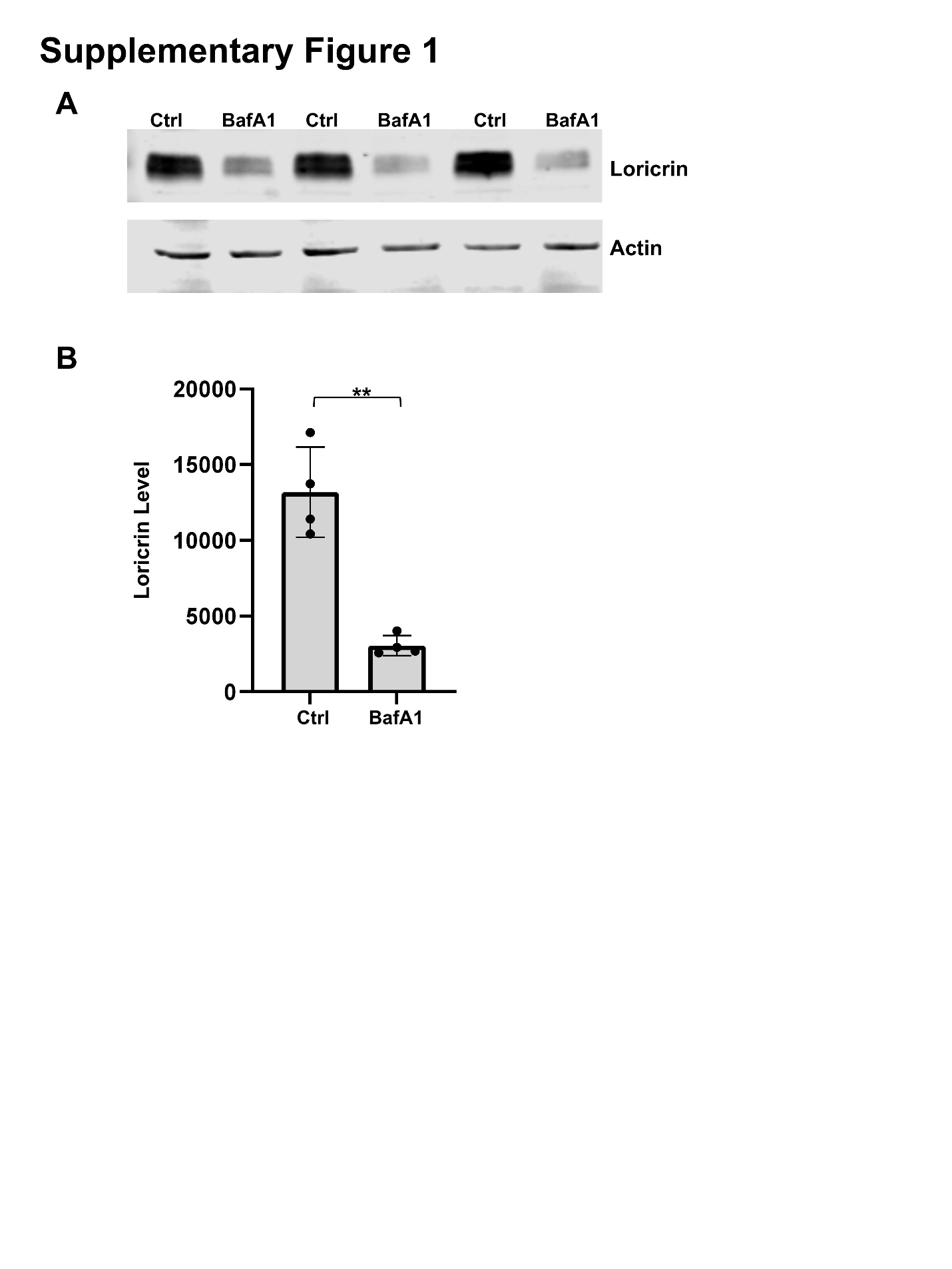

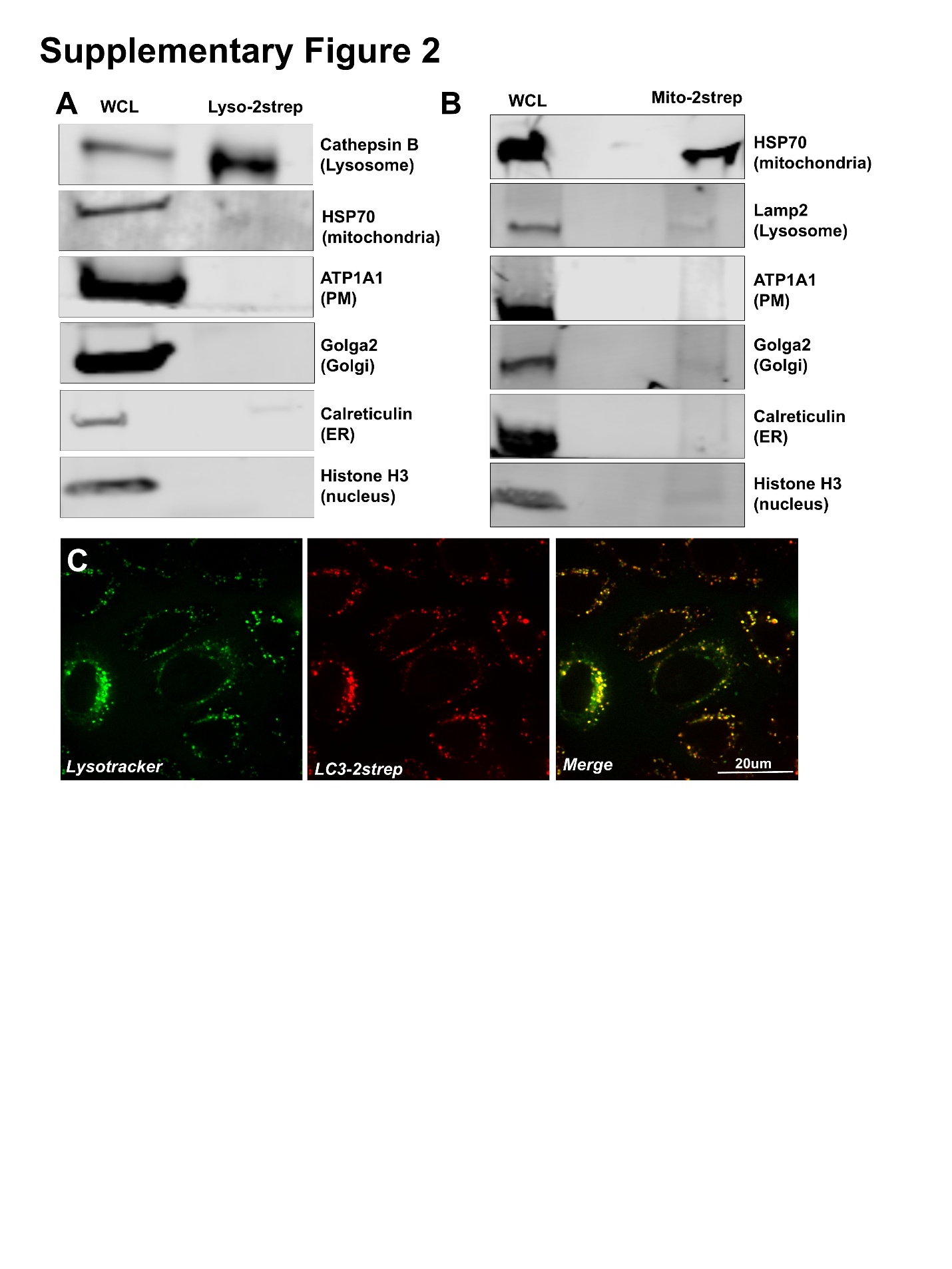

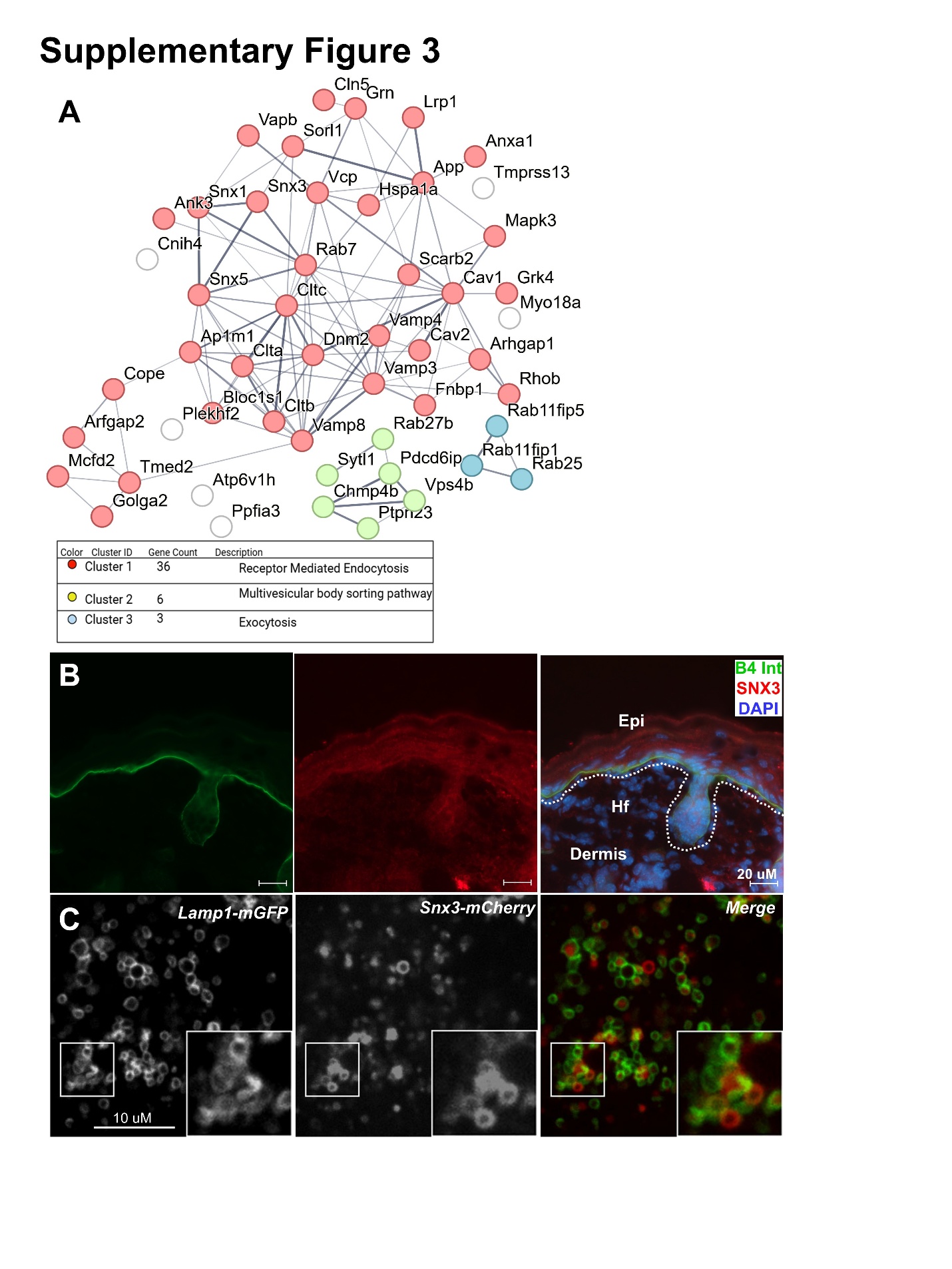

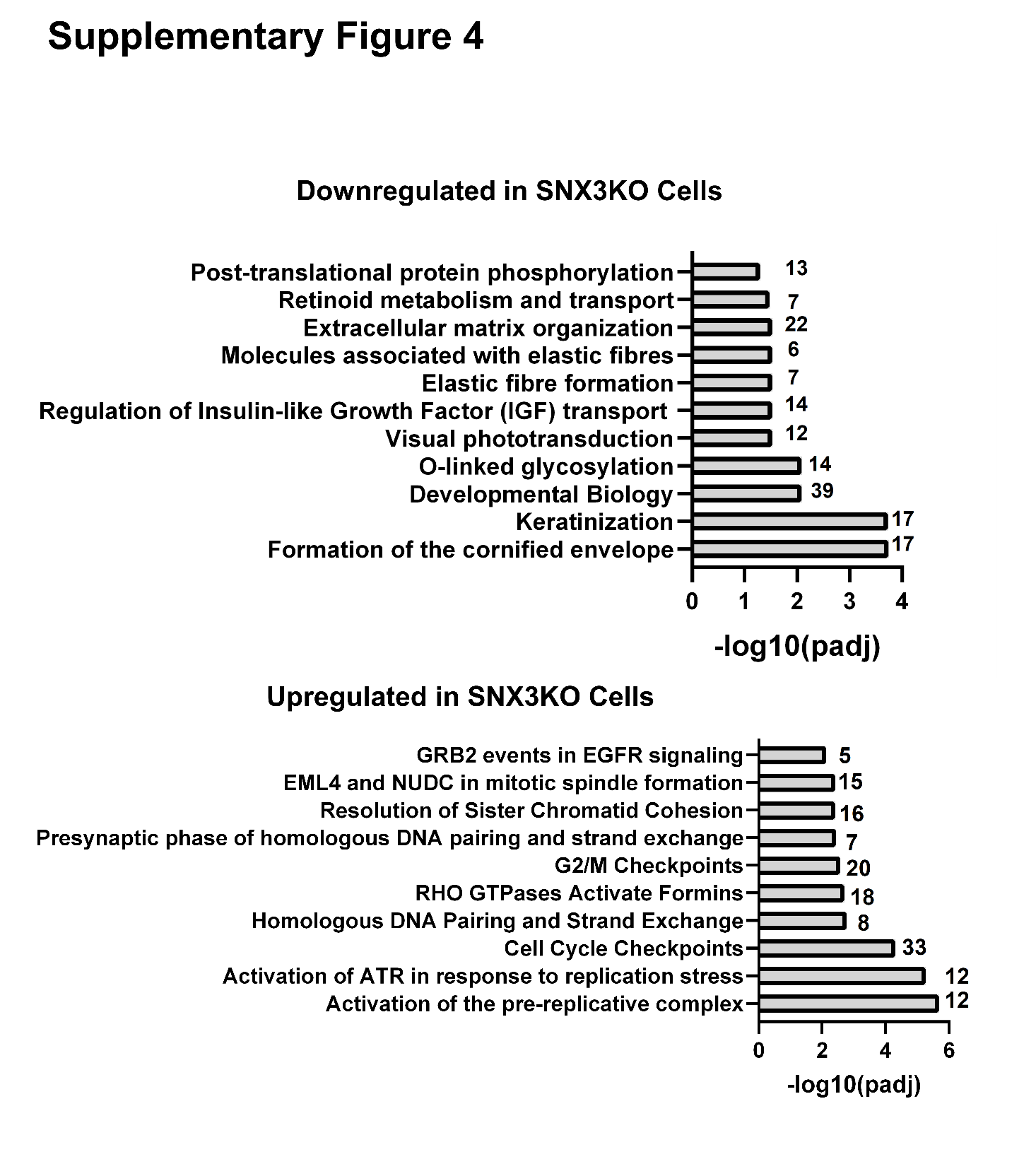

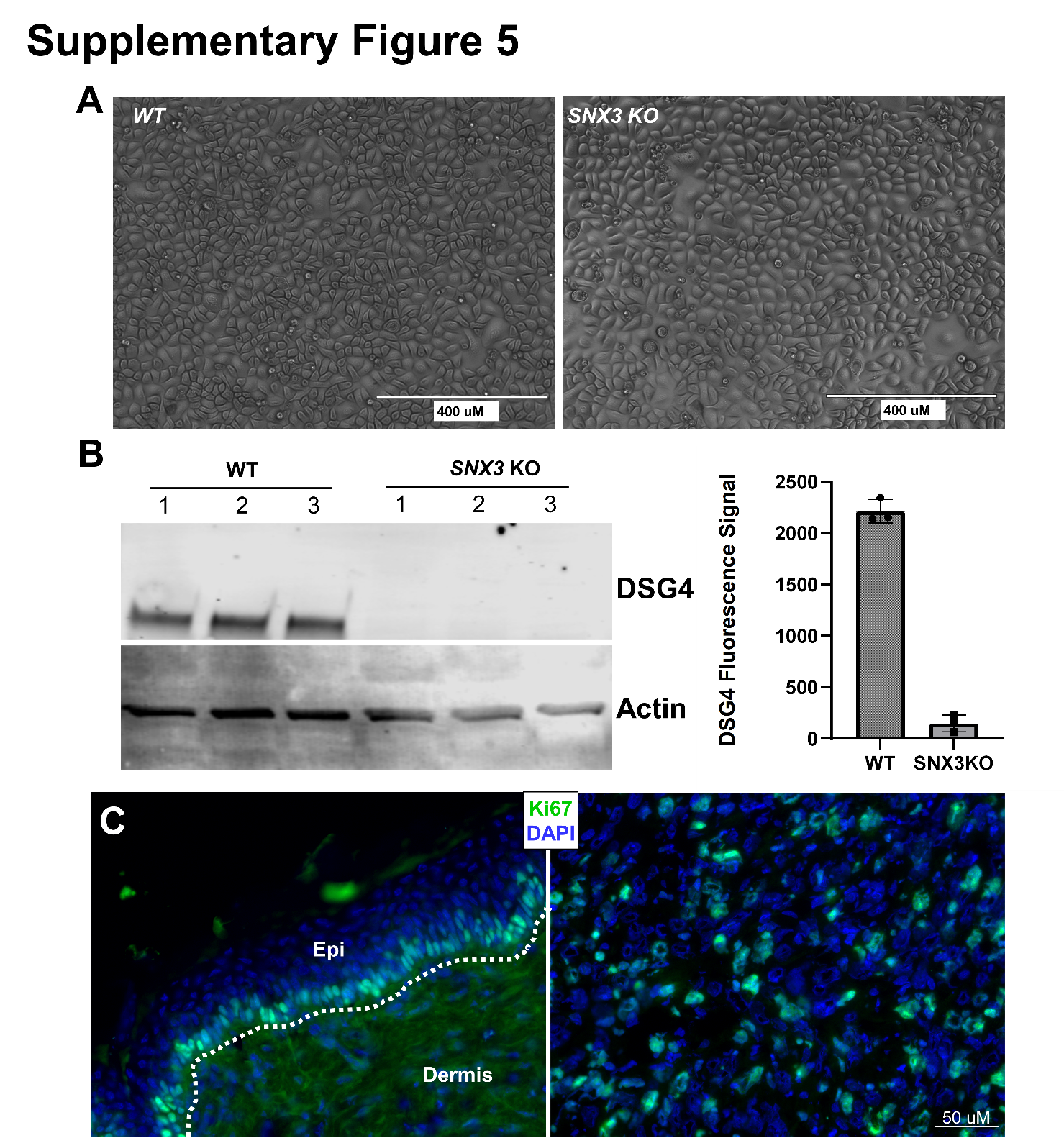

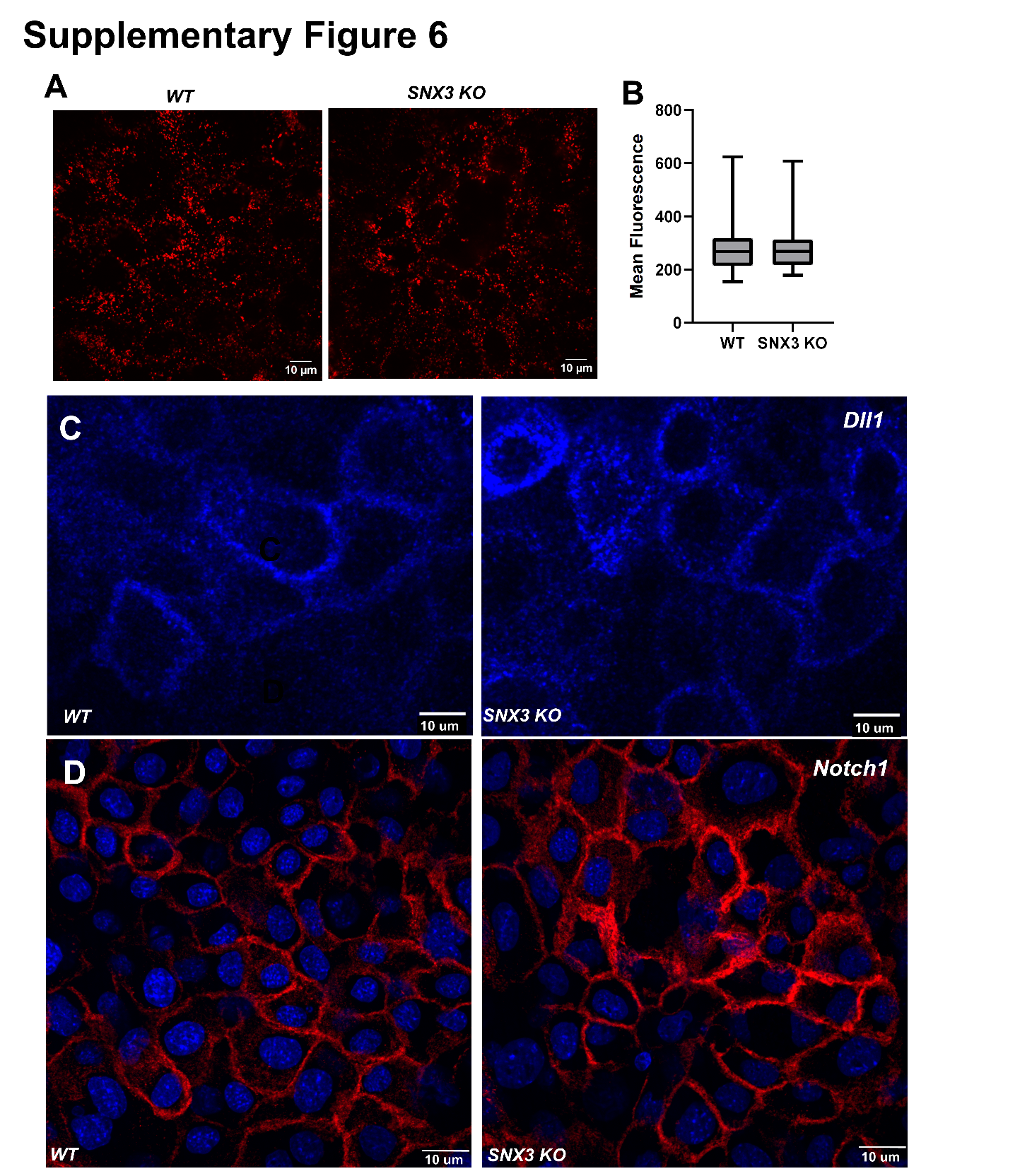

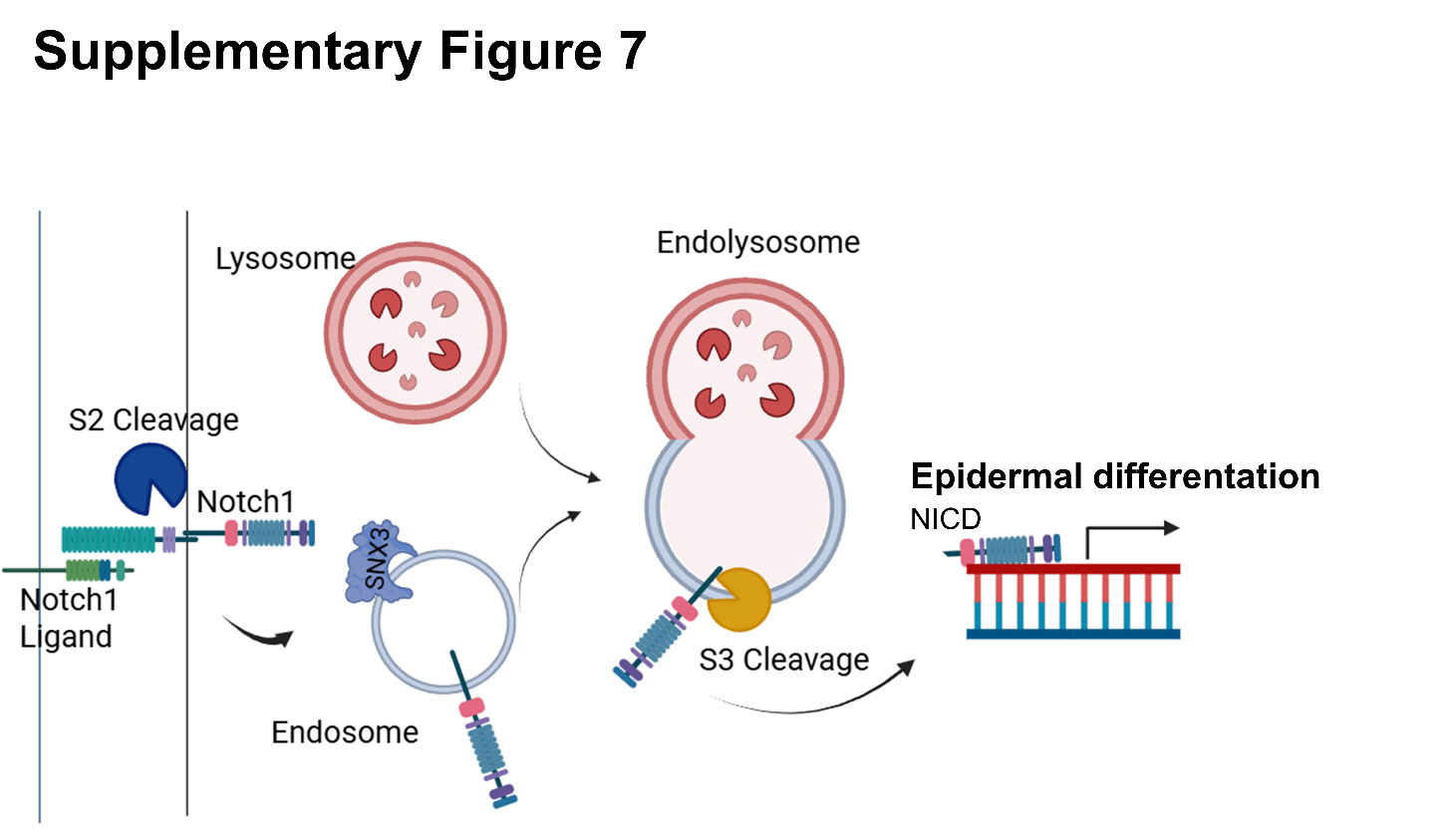
